# Bioengineering of the implantable vascularized endocrine constructs for insulin delivery suitable for clinical upscaling

**DOI:** 10.1101/2025.04.19.647461

**Authors:** Kevin Bellofatto, Fanny Lebreton, Masoud Hassany, Reine Hanna, Juliette Bignard, Antoine Marteyn, Laura Mar Fonseca, Francesco Campo, Cristina Olgasi, Lelia Wolf-van Bürck, Mohsen Honarpisheh, Begoña Martinez de Tejada, Antonia Follenzi, Antonio Citro, Lorenzo Piemonti, Olivier Thaunat, Jochen Seissler, Phillippe Compagnon, Marie Cohen, Ekaterine Berishvili, VANGUARD consortium

## Abstract

Beta cell replacement therapy for type 1 diabetes is hindered by poor graft survival and suboptimal function, largely due to inadequate vascularization and lack of supportive microenvironment. To address these challenges, we developed a clinically scalable, extracellular matrix (ECM)–mimetic hydrogel, termed Amniogel, derived from human amniotic membrane via streamlined, clinically compliant process. Co-encapsulation of pancreatic islets with blood outgrowth endothelial cells (BOECs) within Amniogel facilitated the formation of prevascularized endocrine constructs (VECs). These constructs demonstrated enhanced β-cell viability and function through ECM-bound pro-survival signals, rapid self-assembly of perfusable endothelial networks enabling efficient glucose sensing, and deposition of laminin-rich basement membranes enhancing β-cell coupling and insulin secretion kinetics. In preclinical diabetic mouse models, VECs rapidly integrated with the host vasculature and provided sustained glycemic control when implanted subcutaneously. This integrative approach, combining a scalable, cost-effective biological scaffold with autologous vascularization potential, represents a significant advancement toward durable and clinically translatable β-cell replacement therapies for T1DM.

**One Sentence Summary:** A clinically scalable, biological hydrogel based vascularized endocrine constructs show sustained diabetes reversal.

## INTRODUCTION

Type 1 diabetes mellitus (T1DM) results from autoimmune destruction of insulin-producing β-cells, and while insulin therapy is the standard treatment, it does not prevent long-term complications or ease the daily management burdens (*1*). Intrahepatic islet transplantation temporarily restores insulin independence and improves glycemic control (*2*). However, its widespread clinical implementation is hindered by donor islet scarcity and the risks associated with lifelong immunosuppression (*3–6*). Moreover, instant blood-mediated inflammatory reaction (IBMIR), oxidative stress, loss of extracellular matrix (ECM) signals and poor vascularization significantly compromise islet engraftment and function in the liver (*7–10*). Extrahepatic transplantation sites are being explored to overcome these issues (*11–19*). Amongst these, the subcutaneous space is gaining interest due to its safety, large surface area, and accessibility for implantation and monitoring. However, inadequate vascularization and hypoxia frequently lead to islet graft failure (*20, 21*).

Tissue-engineering approaches aim to mitigate these issues using supportive scaffolds or encapsulation devices. Diverse macrodevices made of synthetic polymers have been designed to house islets or stem cell–derived islet-like clusters (*22*). Although some have advanced to first-in-human trials, clinical success has been limited due to fibrosis, foreign body response, and inadequate vascularization (*23–25*). Critically, macro-devices designed so far rely on inorganic or synthetic polymers, which fail to deliver the native ECM cues and vascularization essential for β-cell survival and function.

Islets are abundantly vascularized miniorgans surrounded by specialized extracellular matrix essential for β-cell viability and proper function (*26*). The isolation process disrupts this irreversibly disrupts this dynamic microenvironment, triggering β-cell apoptosis (*27*). Integrating ECM components with endothelial cells has been shown to improve islet survival and function in preclinical models (*28–32*). Yet, the clinical translation of these strategies remains challenging. An ideal construct would combine a biocompatible biological ECM with a built-in, high-density, functional vascular network in direct contact with encapsulated islets, enabling prompt revascularization, efficient glucose sensing and insulin secretion, and be readily scalable for clinical translation.

Here, we report a clinically scalable strategy to engineer vascularized endocrine constructs for subcutaneous implantation. We first generated Amniogel, an ECM-mimetic hydrogel derived from the human amniotic membrane, suitable for low-cost, clinical-grade manufacturing. By co-encapsulating pancreatic islets and blood outgrowth endothelial cells (BOECs) within this biologic platform, we engineered prevascularized endocrine constructs that replicate key aspects of the native islet niche through three synergistic mechanisms: 1) preservation of β-cell viability and function via essential ECM-bound pro-survival signals, 2) self-assembly of perfusable endothelial networks directly interfacing with islets to enable rapid glucose sensing and 3) deposition of laminin-rich basement membrane components that enhance β-cell gap junction coupling and insulin secretion kinetics. Importantly, engineered constructs demonstrated rapid vascular integration and sustained glycemic control in diabetic mice in two clinically relevant extrahepatic sites—the epididymal fat pad (as an omental analog in rodents) (*33, 34*) and the subcutaneous space. This synergistic strategy, integrating a clinically scalable biomaterial with autologous vascularization potential, represents a promising advance toward clinically translatable, durable cell therapies for type 1 diabetes.

## RESULTS

### Production and characterization of Amniogel

To enable clinical translation of Amniogel, we developed a five-step, Good Manufacturing Practice (GMP)-compliant production protocol (Fig. 1A). Human amniotic membrane (hAM) was decellularized using xenogen-free 10X TrypLE™ Select, effectively reducing DNA content from 424.0 ± 58.15 ng/mg in native tissue to 38.03 ± 2.24 ng/mg, below the biocompatibility threshold of <50 ng/mg (Fig 1B) (*35*). Biochemical profiling showed retention of collagen, although glycosaminoglycan content was partially decreased from 8.83 ± 1.18 to 3.59 ± 1.16 µg/mg (Fig. 1C–D).

**Figure 1.**
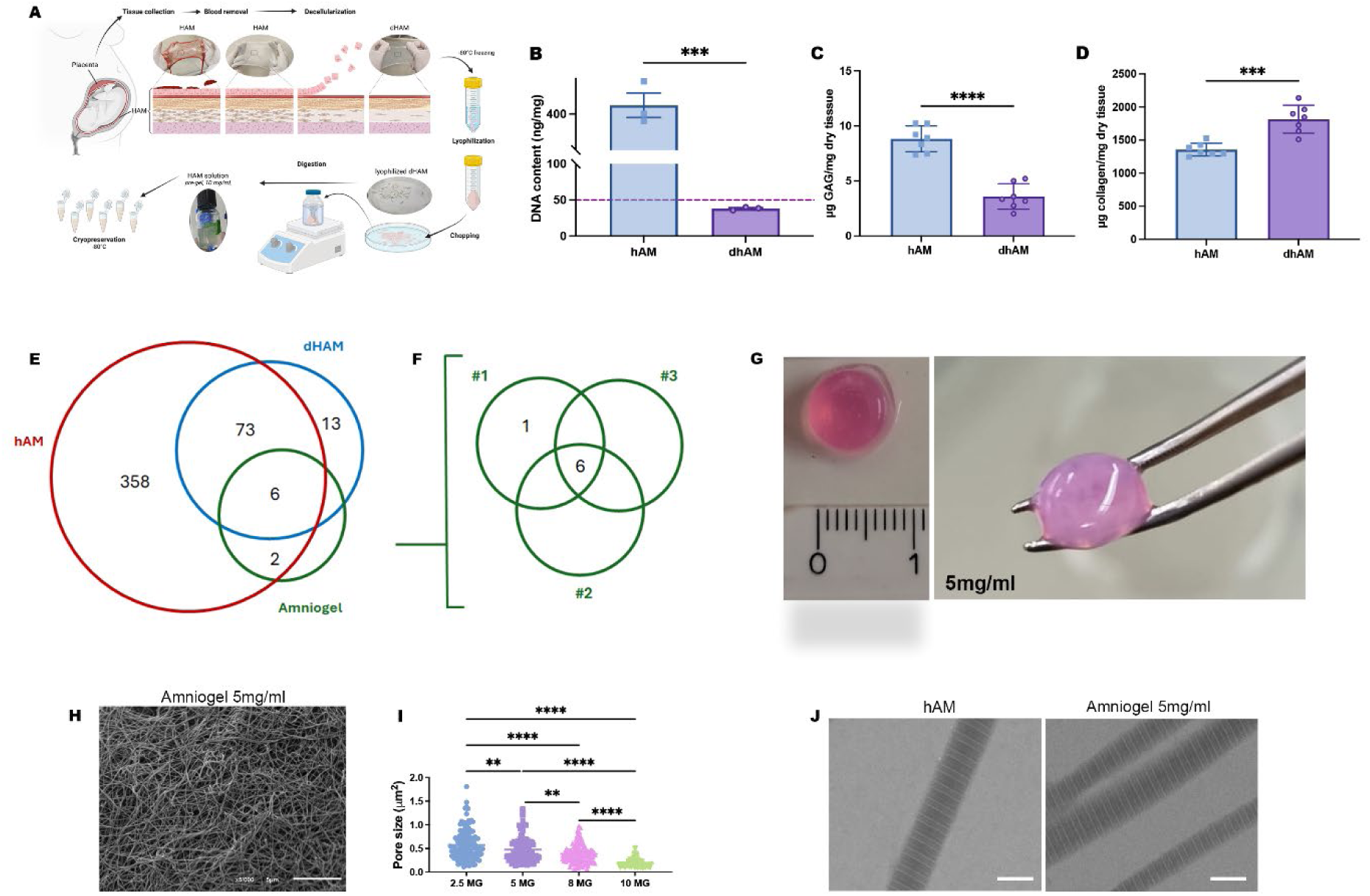
Generation and characterization of Amniogel. (A) Schematic representation of Amniogel preparation. (B) DNA quantification in fresh (hAM) and decellularized (dHAM) human amniotic membranes. Data are presented as mean ± SD, two-tailed unpaired t-test, ***p = 0.0003 (n = 3 biological replicates). (C, D) Quantification of collagen, and GAGs. Data are presented as mean ± SD, two-tailed unpaired t-test, ****p <0.0001, ***p = 0.0002 (n = 7 biological replicates). (E, F) Comparative analysis of ECM protein retention across fresh hAM, dHAM and Amniogel. Venn diagram highlights 92 proteins retained post-decellularization, with six proteins shared across all conditions, including Collagen alpha-1(I), Collagen alpha-2(I), Collagen alpha-3(VI), and Fibrillin-1. The reproducibility of Amniogel protein composition was confirmed across three independent batches, with six proteins consistently retained across all batches and one protein identified in individual batches. (G) Macroscopic view of Amniogel at 5 mg/ml. (H) SEM images of Amniogel (5 mg/ml) displaying an interconnected fibrous network. Scale bars, 5 µm. (I) Pore size analysis of Amniogel. A progressive reduction in pore size was observed between 2.5 and 5 mg/mL, with a pronounced decrease at 8 and 10 mg/mL. Data are presented as mean ± SD, one-way ANOVA with Tukey’s correction: 2.5 mg/ml vs. 5 mg/ml, **p=0.0025; 2.5 mg/ml vs. 8 mg/ml, ****p<0.0001; 2.5 mg/ml vs. 10 mg/ml,****p <0.0001; 5 mg/ml vs. 8 mg/ml, **p=0.0097; 5 mg/ml vs. 10 mg/ml ****p<0.0001; 8 mg/ml vs. 10 mg/ml, ****p<0.0001. Depicted are the mean values of 138 detected pores from six random areas per sample (n = 3 biological replicates). (J, K) TEM analysis revealed no discernible differences in the fiber structure of collagen within Amniogel compared to native human amniotic membrane (hAM). Scale bars, 500 nm.

Following these initial assessments, we analyzed ECM protein retention post-decellularization using mass spectrometry. Across all analyzed samples—native human amniotic membrane (hAM), decellularized hAM (dAM), and three independent batches of Amniogel—we identified a total of 499 proteins. After removing contaminants, sample preparation enzymes, and proteins with a false discovery rate (FDR) below 95%, 452 proteins remained for analysis (Fig. 1E). We then filtered this dataset to focus on ECM proteins known to support islet survival and function (*36*). Among these six were consistently retained in all three Amniogel batches, including collagen alpha-1(I) chain, collagen alpha-1(III) chain, collagen alpha-2(I) chain, collagen alpha-3(VI) chain, and fibrillin-1. Notably, essential cell-binding proteins such as fibronectin were detected, although at reduced abundance. All three independently produced Amniogel batches exhibited highly similar ECM protein profiles, demonstrating robust batch-to-batch reproducibility (Fig.1F).

Amniogels were formulated with ECM concentrations of 2.5–10 mg/ml. Formulations at 5– 10 mg/mL exhibited enhanced structural rigidity, facilitating easy handling and manipulation with forceps (Fig.1G). SEM analysis of Amniogels revealed an inverse relationship between ECM concentration in hydrogel and pore size (0.5712 ± 0.3099 µm at 2.5 mg/mL vs 0.1845 ± 0.08926 µm at 10 mg/mL; Fig. 1H, I). The 5 mg/mL concentration was selected for subsequent experiments due to its optimal balance between handling properties and a pore size that favors the diffusion of oxygen and nutrients. TEM further confirmed that collagen fiber integrity was preserved compared to native hAM (Fig. 1J).

Gelation kinetics of Amniogel were comparable to those of rat tail collagen type I (Fig. S1). Amniogel demonstrated robust cytocompatibility and the capacity for cell-driven remodeling. Our observations show that Amniogel contraction is influenced by both the concentration of decellularized hAM and the cell seeding density. Specifically, when seeded with BOECs, gels with lower stiffness (5 mg/mL) contracted to 37% of their initial area, while stiffer 8 mg/mL gels retained 96% of their original dimensions (Fig. S2A, B). Over seven days, BOECs in 5 mg/mL gels progressively reorganized into interconnected capillary-like networks, whereas 8 mg/mL gels restricted morphogenesis to fragmented, short structures (Fig. S2C). This suggests that Amniogel at 5 mg/mL provides an optimal balance of stiffness, enabling matrix remodeling and supporting reproducible vessel formation.

Furthermore, BOECs cultured on Amniogel-coated plates showed a significant increase in cell numbers relative to collagen controls, adopting spindle-shaped morphologies and proliferating over seven days (Fig. S3A, B). Viability (FDA/PI) assays confirmed minimal cytotoxicity, with >90% cell survival across all conditions, validating Amniogel’s biocompatibility (Fig. S3C). These results demonstrate that Amniogel is a biocompatible scaffold that retains essential ECM components crucial for islet viability and function, while supporting cell attachment and matrix remodeling.

### Islet survival and function in Amniogel

We next evaluated whether Amniogel encapsulation protects and improves islet function. Under normoxic conditions (20% O₂), viability of rat islets embedded in Amniogel was similar to controls cultured in suspension (Fig. S4A, C, E). However, suspension-cultured islets formed clumps and developed necrotic cores over time. Under hypoxic conditions (1% O₂ for 16 h), suspended islets showed extensive necrosis, while Amniogel-embedded islets maintained high viability (Fig. S4B to E). Notably, Amniogel-embedded islets demonstrated significantly enhanced glucose-stimulated insulin secretion and a higher stimulation index (SI: 0.7 ± 0.3 vs 2.3 ± 0.95, p = 0.05) compared to hypoxic controls (Fig. S4F). These results suggest that Amniogel attenuates hypoxic injury and preserves islet functionality under stress.

To evaluate the impact of cellular adhesion to Amniogel on human β-cell function and viability, human islets were embedded in Amniogel to form endocrine constructs (hEC) or maintained in standard suspension culture (HI alone) as controls for 7 days. Islets embedded in Amniogel retained their normal morphology and hormone expression (Fig. 2A,B). Basal insulin release was reduced in hECs, suggesting improved glucose regulation (Fig. S5A). Furthermore, hECs showed significantly enhanced insulin secretion both under high-glucose stimulation (SI: 2.317 ± 0.195 vs 1.204 ± 1.135, p = 0.0012) and theophylline stimulation (SI: 4.240 ± 0.673 vs 2.385 ± 0.110, p = 0.0092) compared to HI alone (Fig. S5B, C). To elucidate mechanisms underlying enhanced insulin secretion in hECs, we analyzed β1-integrin expression. Immunofluorescence revealed a significantly increased β1-integrin expression in Amniogel-embedded islets compared to controls (Fig. 2C, D). This was confirmed by qPCR, showing a threefold increase in *ITGB1* expression (p= 0.0016; Fig. 2E). The elevated β1-integrin expression was associated with significantly reduced pro-apoptotic *CASP3* expression (p = 0.0003) and increased anti-apoptotic *BCL2* expression by a 4.8-fold (p=0.0089; Fig. 2F, G).

**Figure 2.**
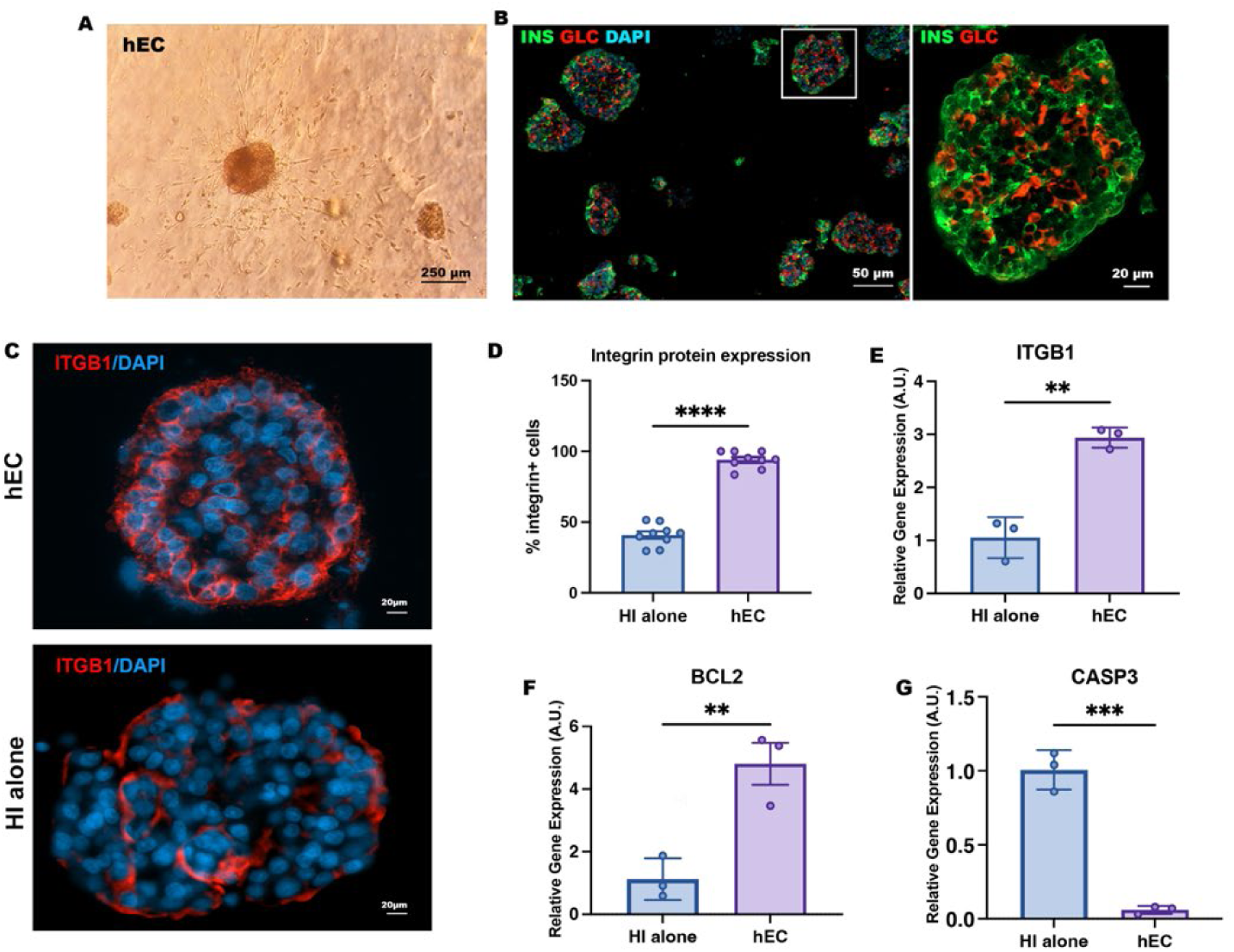
Amniogel Enhances Human Islet Viability and Function. (A, B) Optical microscopy and immunofluorescent images of Amniogel-embedded human islets cultured for 7 days. Islets were stained for insulin (green) and glucagon (red). (C–D) Representative immunofluorescence images and quantification of β1-integrin expression in hECs and human islets in suspension. Data are presented as mean ± SD, two-tailed unpaired t-test, ****p < 0.0001. (E-G) qPCR of *ITGB CASP3* and *BCL2* on hECs and HI alone. Data are presented as arbitrary units (AU) after normalization to housekeeping genes, mean ± SD, two-tailed unpaired t-test, *ITGB,* **p = 0.0016; *BCL2*, **p = 0.0089; *CASP3*, ***p = 0.0003 (n = 3 biological replicates).

These findings demonstrate that Amniogel promotes β-cell viability, structural integrity, and insulin secretion by enhancing cellular interactions and reducing apoptosis.

### Amniogel supports endothelial cell self-organization into a vascular network

To evaluate Amniogel’s ability to support endothelial cell self-organization, LV-GFP BOECs were embedded in Amniogel and cultured in vasculogenic media (VM) (Fig. 3A).

**Figure 3.**
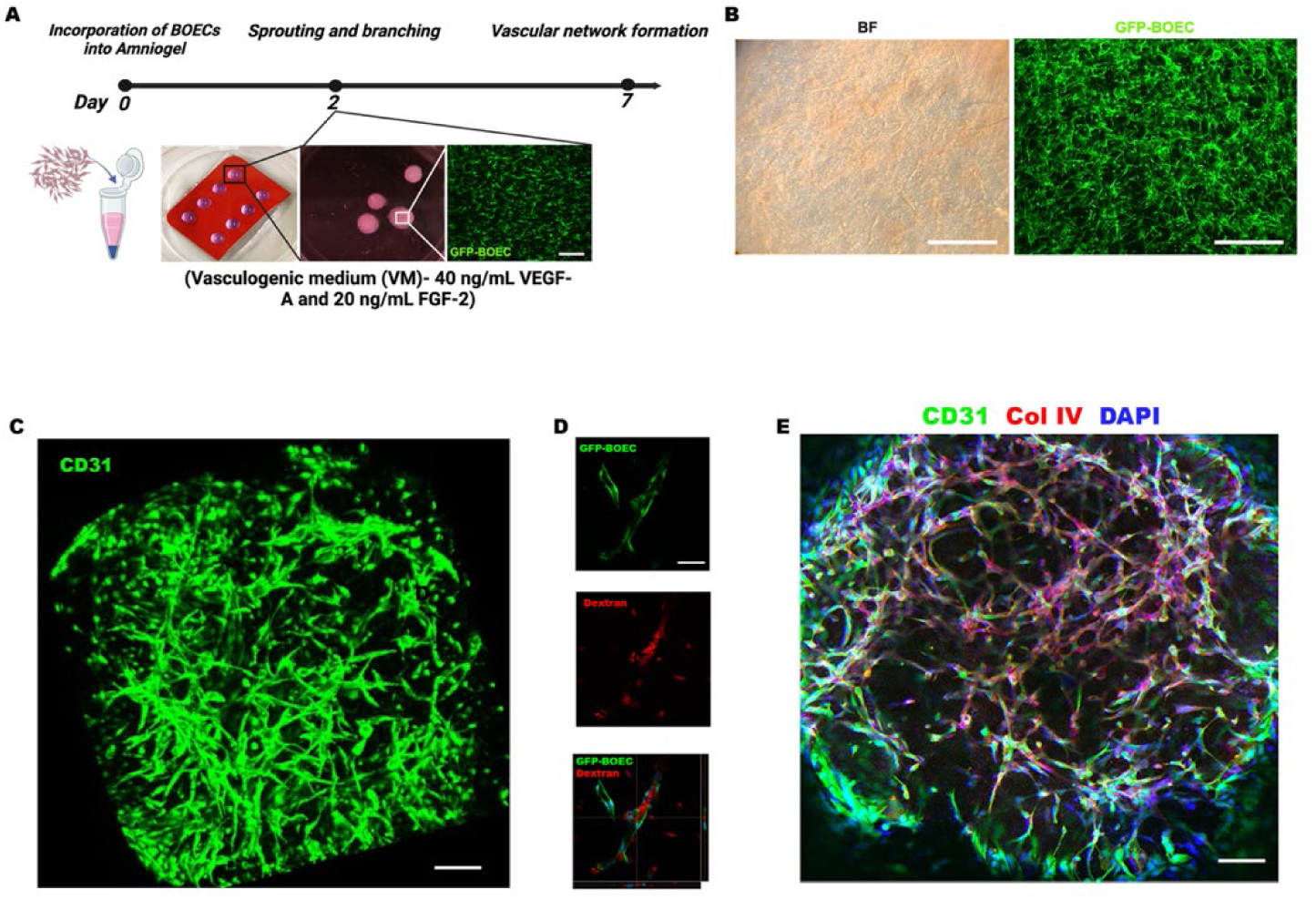
Amniogel Facilitates Formation of vascular networks. (A) Schematic representation of vascular construct generation. Scale bars, 500 μm. Created in https://BioRender.com (B) Representative bright-field and immunofluorescence images of LV-GFP transduced BOECs showing the establishment of vascular networks. Scale bars, 500 μm. (C) 3D reconstruction of capillary organization (CD31, green) in a vascular construct. Scale bars, 50 μm. (D) Formation of hollow endothelial structures within 3D constructs, visualized through virtual stacks of LV-GFP transduced BOECs incubated with Texas Red-labeled dextran. Scale bars, 30 μm. (E) Whole-mount immunofluorescence of vascular networks showing endothelial cells (CD31, green) and the vascular basement membrane (Col IV, red). Scale bars, 50 µm.

Cells formed interconnected structures within 48 hours, evolving into a robust vascular network by day 7 (Fig. 3B). Individual cells were no longer distinguishable, indicating their complete integration into continuous tubular structures (Fig. 3C). Functional validation was demonstrated by Texas red–labeled dextran perfusion into hollow lumens, confirming effective fluid transport (Fig. 3D). Immunostaining further demonstrated that the vessel-like structures were surrounded by a basement membrane, as evidenced by collagen type IV deposition (Fig. 3E). These findings indicate that Amniogel provides a supportive microenvironment facilitating endothelial cell self-organization into functional vascular networks.

### Generation of vascularized endocrine constructs

We hypothesized that Amniogel could facilitate the synergistic interaction between endothelial cells and pancreatic islets, thereby supporting the development of more complex vascularized endocrine constructs (hVECs). To test this hypothesis, we incorporated both human derived islets and BOECs into Amniogel and cultured them for 7 days in optimized vasculogenic medium (OVM) (Fig. 4 A). Within 3 days, BOECs formed continuous 3D vascular networks around islets (Fig. 4B). Interestingly, despite the initial lack of laminin in Amniogel, the islets were surrounded by a laminin-rich matrix (Fig. 4C). qPCR analysis identified BOECs as the primary laminin source, showing a 3.9-fold increase in *LAMB1* expression in hVECs (p=0.008) and 5.7-fold in BOECs (p=0.0003) versus HI alone (Fig. 4D). This indicates that BOECs play a critical role in laminin deposition within the gel, leading to the development of a physiological-like ECM surrounding the islets.

**Figure 4.**
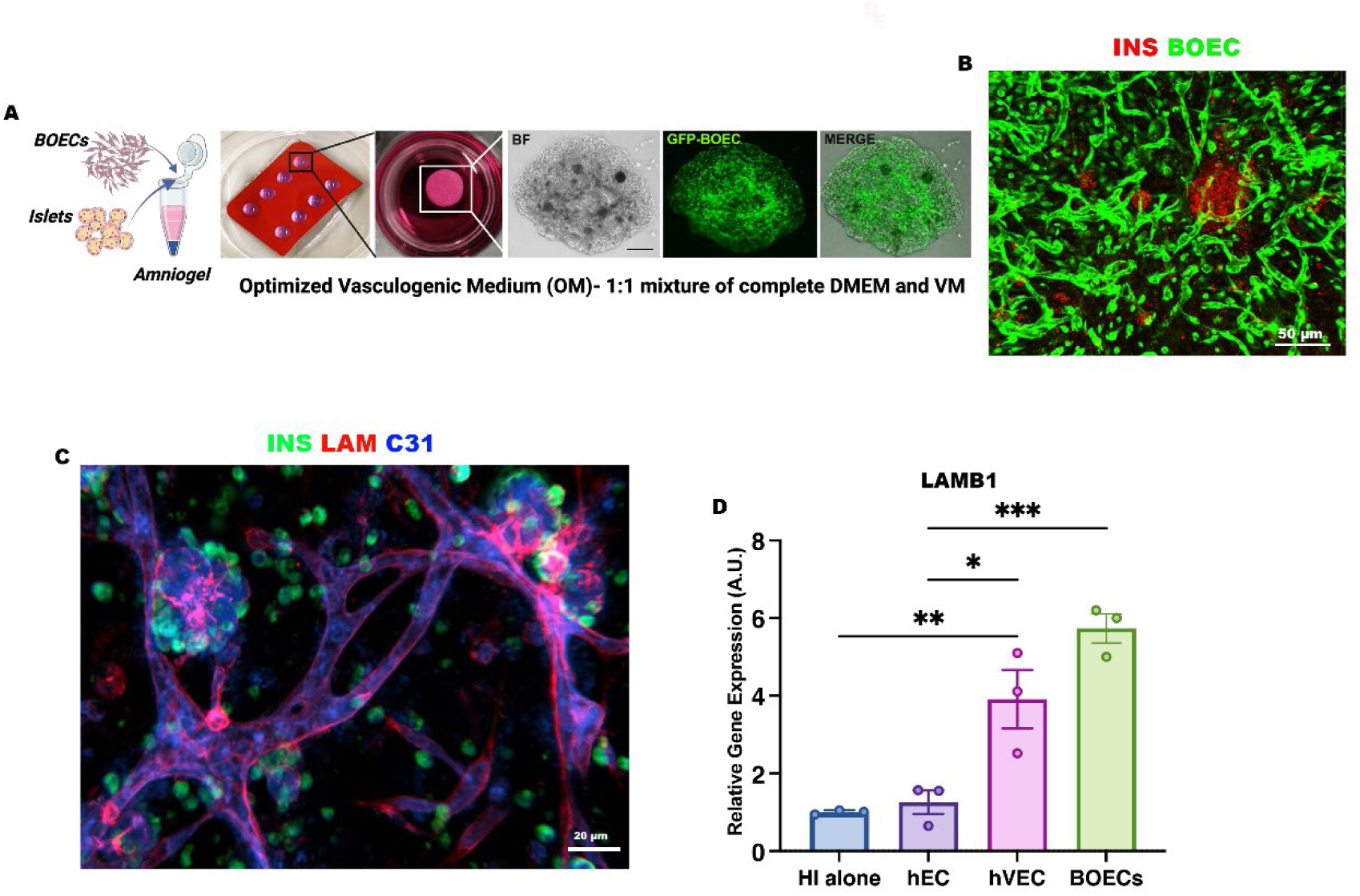
Assembly of hVECs. (A) Schematic representation of hVEC assembly. Representative bright-field and immunofluorescence images of hVECs demonstrate the formation of a dense vascular network (green, GFP-transduced BOECs) surrounding the embedded islets. Scale bar, 200 μm. Created in https://BioRender.com. (B) Representative confocal fluorescence microscopy images of hVEC reveal vessel formation (green, GFP-transduced BOECs) adjacent to islets (insulin, red). Scale bar, 50mm. (C) Immunofluorescent staining of laminin (red) highlights the presence of basement membrane encasing islets (insulin, green) and CD31+ endothelium (blue). Scale bar, 20mm.

To explore functional implications of vascular integration, we investigated the role of gap junctions, specialized membrane domains composed of nonspecific channels that facilitate cell-to-cell communication, which is essential for coordinated insulin release (Fig. 5A) (*37*). qPCR analysis revealed significant upregulation of *GJD2,* with increases observed in both hECs and hVECs (5.3-fold, p = 0.0057, and 8.8-fold, p < 0.0001, respectively) compared to HI alone (Fig. 5B). Similarly, expression of *GJA1* increased notably, rising 6.3-fold in hECs (p = 0.0042) and 11.4-fold in hVECs (p< 0.0001) relative to controls (Fig. 5C), indicating strong coupling between β cells and endothelial cells. Consistent with these molecular findings, functional glucose-stimulated insulin secretion (GSIS) assays showed superior performance in vascularized constructs, characterized by reduced basal secretion and increased glucose responsiveness (Fig. S6). Together, these data suggest engineered vasculature within Amniogel enhances islet function through dual mechanisms: (i) BOEC-derived laminin deposition recapitulating native peri-islet ECM, and (ii) connexin-mediated cell–cell communication essential for coordinated insulin release.

**Figure 5.**
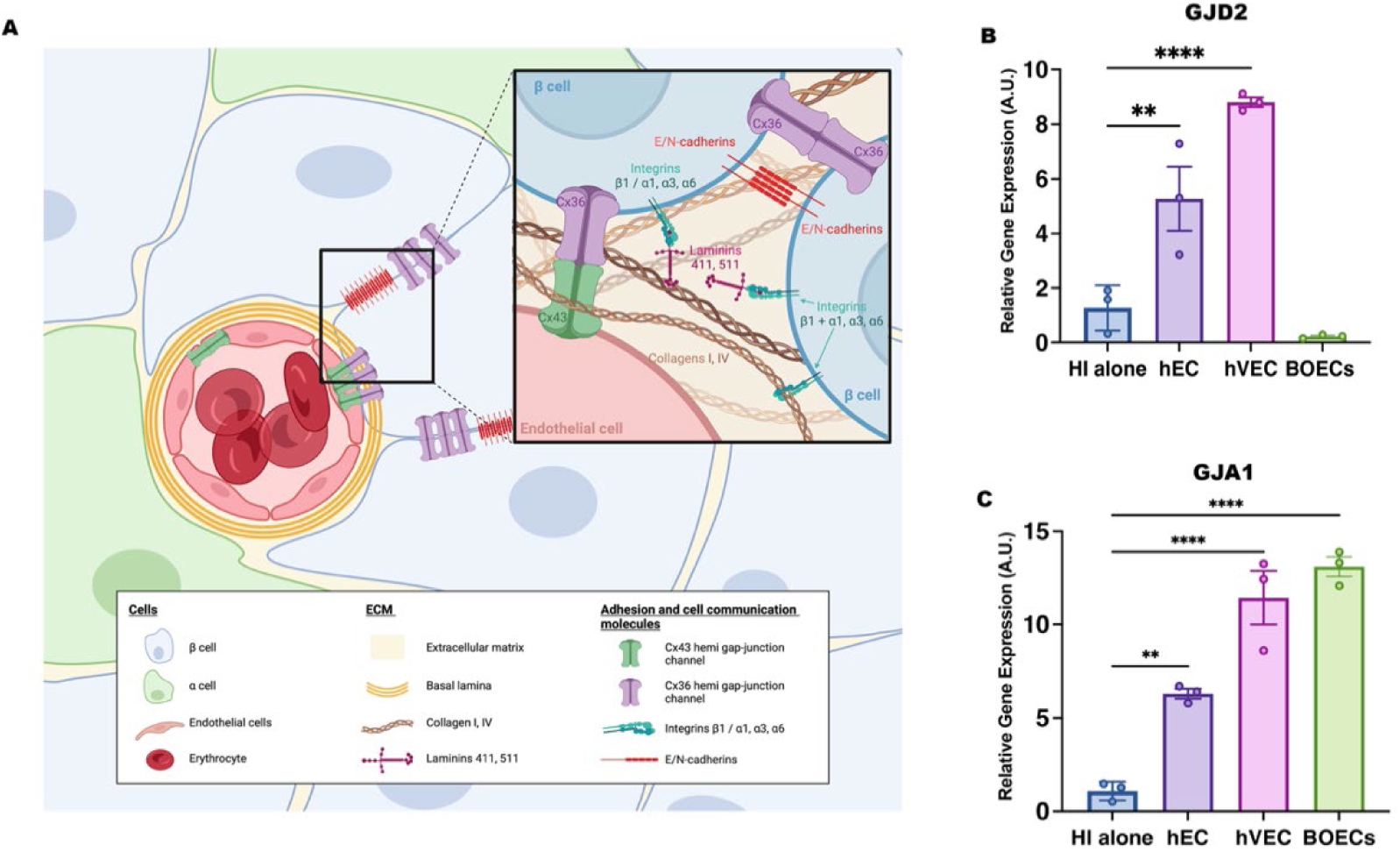
Engineering vasculature within amniogel promotes islet intercellular and cell-to-matrix communication. (A) Graphical illustration of known intercellular and ECM–cell interactions within islets. Cell adhesion molecules enhance β-cell glucose responsiveness, while connexin 36 and connexins 36/43 mediate insulin release by coordinating communication between β-cells and endothelial cells. Cadherins support cell–cell adhesion and inhibit apoptosis in α and β-cells. The ECM, enriched with laminin-211, laminin-511, and type IV collagen, forms a microenvironment that promotes granule secretion, cell proliferation, and apoptosis inhibition. Integrins bind ECM components like collagen and laminin, activating signaling cascades that regulate cell survival, growth, and differentiation. Created in https://BioRender.com (B, C) qPCR of *GJD2 and GJA1* in hECs, hVECs and HI alone. Data are presented as arbitrary units (AU), normalized to housekeeping genes, and shown as mean ± SD. One-way ANOVA with Tukey’s correction, *GJD2:* hEC vs HI alone, **p = 0.0057; hVEC vs HI alone, ****p<0.0001; HI alone vs BOECs, n.s. p= 0.5452 (n = 3 biological replicates); *GJA1*: hEC vs HI alone, **p = 0.0042; hVEC vs HI alone, **** p<0.0001; HI alone vs BOECs, **** p<0.0001 (n = 3 biological replicates).

### In vivo biocompatibility

To determine whether Amniogel triggers an immune response upon transplantation, empty hydrogels were transplanted subcutaneously into immunocompetent C57BL/6 mice (Fig. S7A). Grafts were harvested 1month post-implantation and analyzed histologically. H&E staining revealed modest mononuclear cell infiltration at the implantation sites without granuloma formation, indicating minimal local toxicity and high tissue compatibility (Fig. S7B). Masson’s trichrome staining demonstrated preserved collagen fibrils in the Amniogel. Spindle-shaped cells, likely fibroblasts, were also observed within the grafts. Immunohistochemical staining for CD45 and CD11b confirmed that immune cell infiltration in transplanted acellular gels was comparable to that in sham-operated controls (Fig. S7B).

### Site-specific engraftment of non-vascularized endocrine constructs

We then tested the performance of the Amniogel-based constructs in vivo using streptozotocin-diabetic NSG immunodeficient mice (Fig. 6A). In the highly vascularized epididymal fat pad (EFP) site, even non-vascularized endocrine constructs (EFP-rEC) significantly outperformed islet transplanted alone (EFP-rI). All mice receiving EFP-rECs achieved normoglycemia within one week, whereas only ∼55% in the EFP-rI group became normoglycemic (Fig. 6B, C). An intraperitoneal glucose tolerance test (IPGTT) at 5 weeks post-transplant showed that EFP-rEC recipients cleared glucose almost as efficiently as healthy nondiabetic mice, whereas EFP-rI mice displayed impaired glucose tolerance (Fig. 6D, E). Removing the graft-bearing EFP led to recurrence of hyperglycemia in cured mice, confirming that the implanted constructs were responsible for diabetes reversal. Histological analysis at 14 weeks showed well-preserved islets in the EFP-rEC grafts, with a larger insulin-positive area compared to islet-alone grafts (Fig. 6F, G). Immunohistochemical staining for CD31 revealed a markedly higher vessel density in EFP-rEC samples compared to controls (Fig. 6F, H), suggesting enhanced vascularization and engraftment.

**Figure 6.**
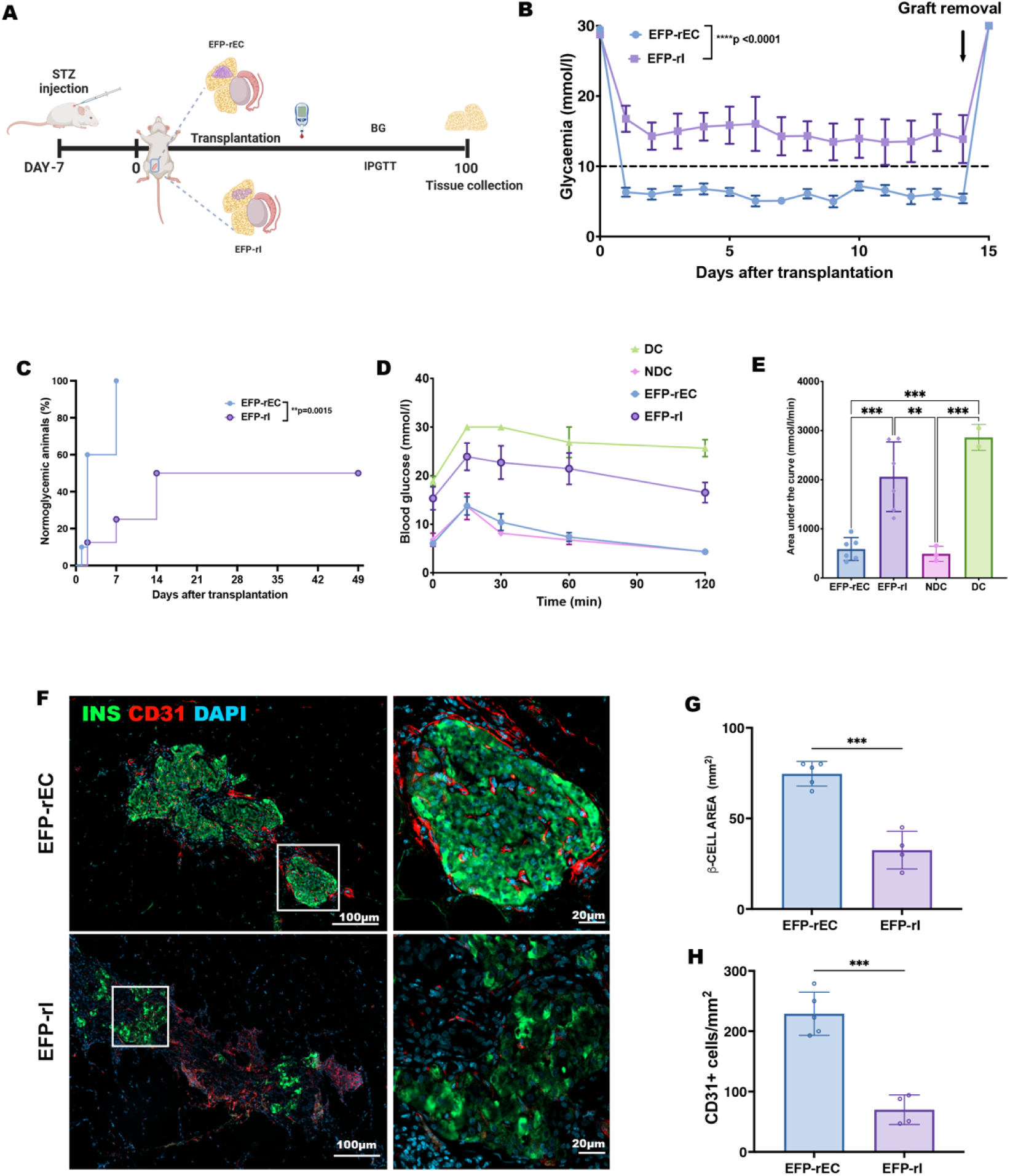
Function of endocrine constructs transplanted in EFP. (A) Schematic representation of the experimental setup. (B) Blood glucose measurements following transplantation. Data are presented as mean ± SEM, analyzed by two-way ANOVA with Sidak correction, ****p < 0.0001. (C) Kaplan-Meier analysis of the percentage of mice achieving normoglycemia (≤11 mmol/L). Differences between EFP-rEC and EFP-rI groups were assessed using a two-sided log-rank (Mantel–Cox) test, **p = 0.0015. (D) Blood glucose curves from the IPGTT on day 30 for EFP-rEC (n = 6), EFP-rI (n = 6), non-diabetic healthy controls (NDC, n = 3), and diabetic controls (DC, n = 2). Data are presented as mean ± SEM. (E) AUC analysis of IPGTT on day 30 for EFP-rEC (n = 6), EFP-rI (n = 6), NDC (n = 3), and DC (n = 2). Data are presented as mean ± SD, analyzed by one-way ANOVA with Tukey’s correction: EFP-rEC vs. EFP-rI, ***p = 0.0006; EFP-rEC vs. DC, ***p = 0.0003; EFP-rI vs. NDC, **p = 0.0021. (F) Immunohistochemical staining of retrieved constructs on day 100 showing insulin (green) expression and vascularization (CD31, red). (F) Representative immunofluorescent images of rat insulin (green) and mouse blood vessels (CD31, red) in a retrieved constructs on day 100. (G) Percentage of insulin-positive area in EFP-rEC (n=5) and EFP-rI (n=4) visualized 100 days after transplantation. Data are presented as mean ± SD, two-tailed unpaired t-test, ***p=0.0002 (H) Blood vessel density in the EFP-rEC (n=5) and EFP-rI (n=4); Data are presented as mean ± SD, two-tailed unpaired t-test, ***p=0.0001.

Despite these encouraging results, the subsequent subcutaneous transplantation of non-vascularized endocrine constructs (SQ-rEC) proved unsuccessful. None of the recipients achieved normoglycemia, demonstrating that simply providing an ECM scaffold was insufficient to support islet function in this poorly vascularized environment (Fig. S8). These findings confirmed that prevascularization is essential for graft survival and efficacy in poorly vascularized subcutaneous site.

### Vascularized endocrine constructs improve glycemic control in subcutaneous transplants

Building on *in vitro* findings demonstrating Amniogel’s capacity to support vascular network formation and enhance islet function through intercellular and cell-to-matrix communication, we moved forward to evaluate the in vivo functionality of VECs in a clinically relevant but poorly vascularized subcutaneous site.

To this end, the vascularized constructs (SQ-rVEC), were implanted subcutaneously in diabetic NSG mice. For comparison, two additional groups of mice were transplanted with non-vascularized constructs containing islets (SQ-rEC) or with islets alone (SQ-rI). Graft function was monitored for 14 weeks (Fig. 7A). Remarkably, 96% of SQ-rVEC recipients achieved rapid normoglycemia within one week, maintaining stable glucose levels significantly lower than both control groups (p<0.001) (Fig. 7B, C). Glucose tolerance tests at day 30 showed that SQ-rVEC group responded to the glucose challenge similarly to healthy controls and significantly better than the SQ-rEC and SQ-rI groups (Fig. 7D, E).

**Figure 7.**
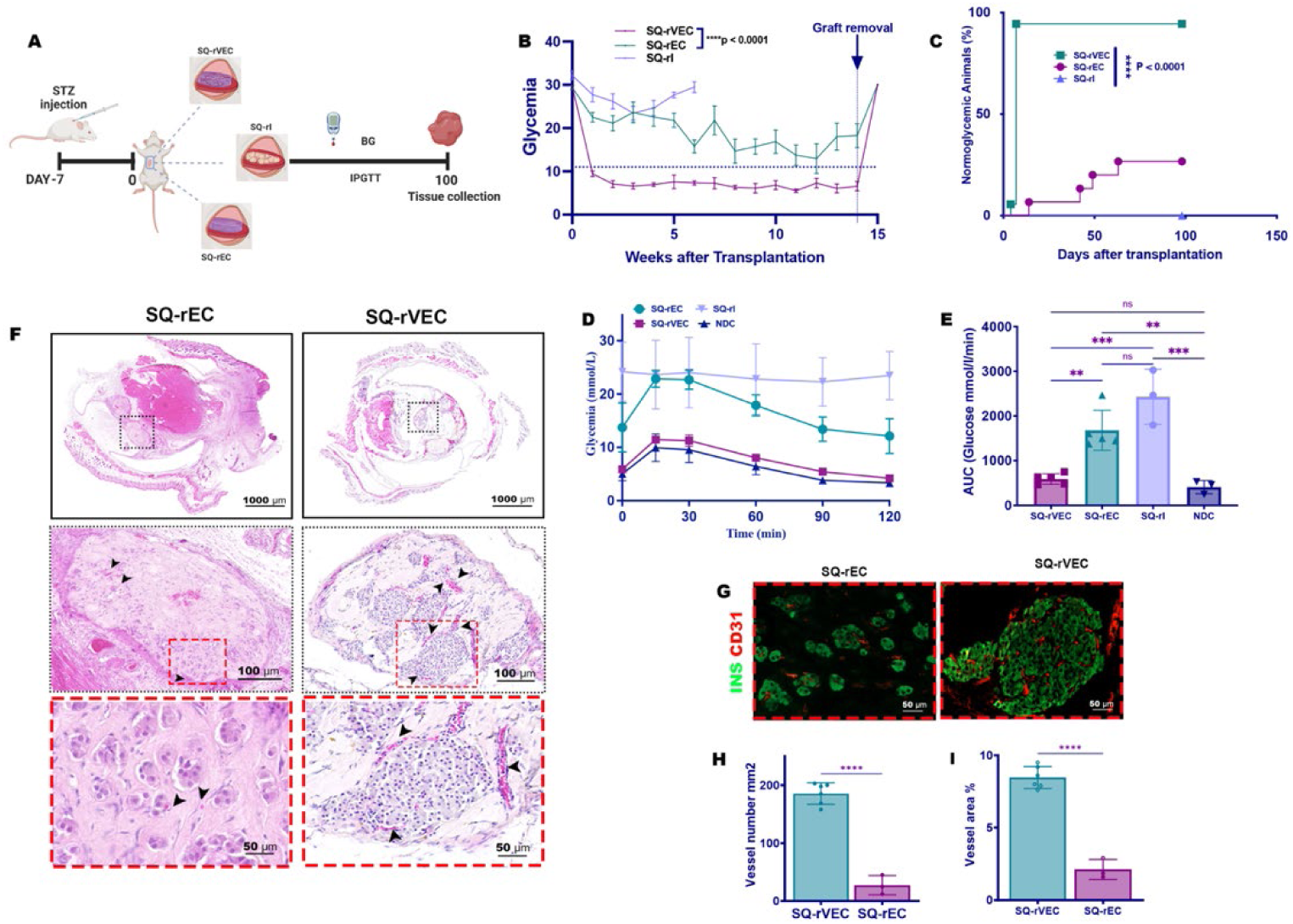
Performance of rat derived VECs after subcutaneous transplantation in diabetic mice. (A) Schematic representation of the experimental setup. (B) Blood glucose measurements following transplantation. Data are presented as mean ± SEM, analyzed by two-way ANOVA with Sidak correction, ****p < 0.0001. (C) Kaplan-Meier analysis of the percentage of mice achieving normoglycemia (≤11 mmol/L). Differences between SQ-rVEC, SQ-rEC and SQ-rI groups were assessed using a two-sided log-rank (Mantel–Cox) test, ****p = 0.0001. (D) Graft function evaluated by IPGTT on day 30 for SQ-rVEC (n = 5), SQ-rVEC (n = 5), SQ-rI (n = 3) and NDC (n = 3). Data are presented as mean ± SEM (E) AUC analysis of IPGTT. Data are presented as mean ± SD, one-way ANOVA with Tukey’s correction, SQ-rVEC v.s. SQ-rI, ***p=0.0001, SQ-rVEC vs. SQ-rEC, **p=0.0028; SQ-rVEC vs. NDC, n.s. 0.9101; SQ-rI vs. SQ-rEC, n.s. p=0.0700; SQ-rI vs. NDC, ***p=0.0001; SQ-rEC vs. NDC, **p=0.0026; (F) H&E-staining of retrieved constructs on day 100. Black arrowheads indicate blood vessels containing RBCs; (G) Immunohistochemical staining of retrieved constructs on day 100 showing insulin (green) expression and vascularization (CD31, red). (H, I) Blood vessel density and area percentage in the SQ-rVEC (n=6) and SQ-rEC (n=3); Data are presented as mean ± SD, two-tailed unpaired t-test, ****p<0.0001.

Graft excision led to diabetes recurrence, confirming graft function. Histological analysis revealed well-vascularized, healthy islets in SQ-rVECs, contrasting with the smaller, fragmented, and poorly vascularized islets in SQ-rECs (Figure 7G). Notably, the implanted matrix remained intact and was repopulated by host cells in both groups. Quantification of blood vessels within and surrounding the constructs (Figure 7H).

To evaluate clinical potential, we assessed subcutaneous transplantation of human islets. Given mice’s inherent resistance to human insulin and the requirement for a larger islet mass to achieve normoglycemia in xenotransplantation studies (*38–44*), two islet-laden vascularized constructs (SQ-hVEC) were implanted bilaterally into the ventral subcutaneous space. Control mice received equivalent non-vascularized endocrine constructs (SQ-hEC). To minimize graft loss resulting from innate immune responses known to occur even in severely immunodeficient mice (*45–47*), experiments were limited to 45 days. This timeframe is clinically relevant, as islet function at one month, assessed by the BETA-2 score (*48*), reliably predicts long-term transplant success (*49*).

SQ-hVEC group demonstrated superior outcomes, with 89% (8/9) achieving normoglycemia within 14 days versus 33% (2/6) in SQ-hEC group (Fig. 8A, B). Human C-peptide levels correlated with glycemic control (Fig. 8C), and glucose tolerance tests showed significantly improved glucose clearance in SQ-hVEC mice (p=0.0057, Fig. 8D). All reverted to hyperglycemia upon graft removal, confirming therapeutic dependence.

**Figure 8.**
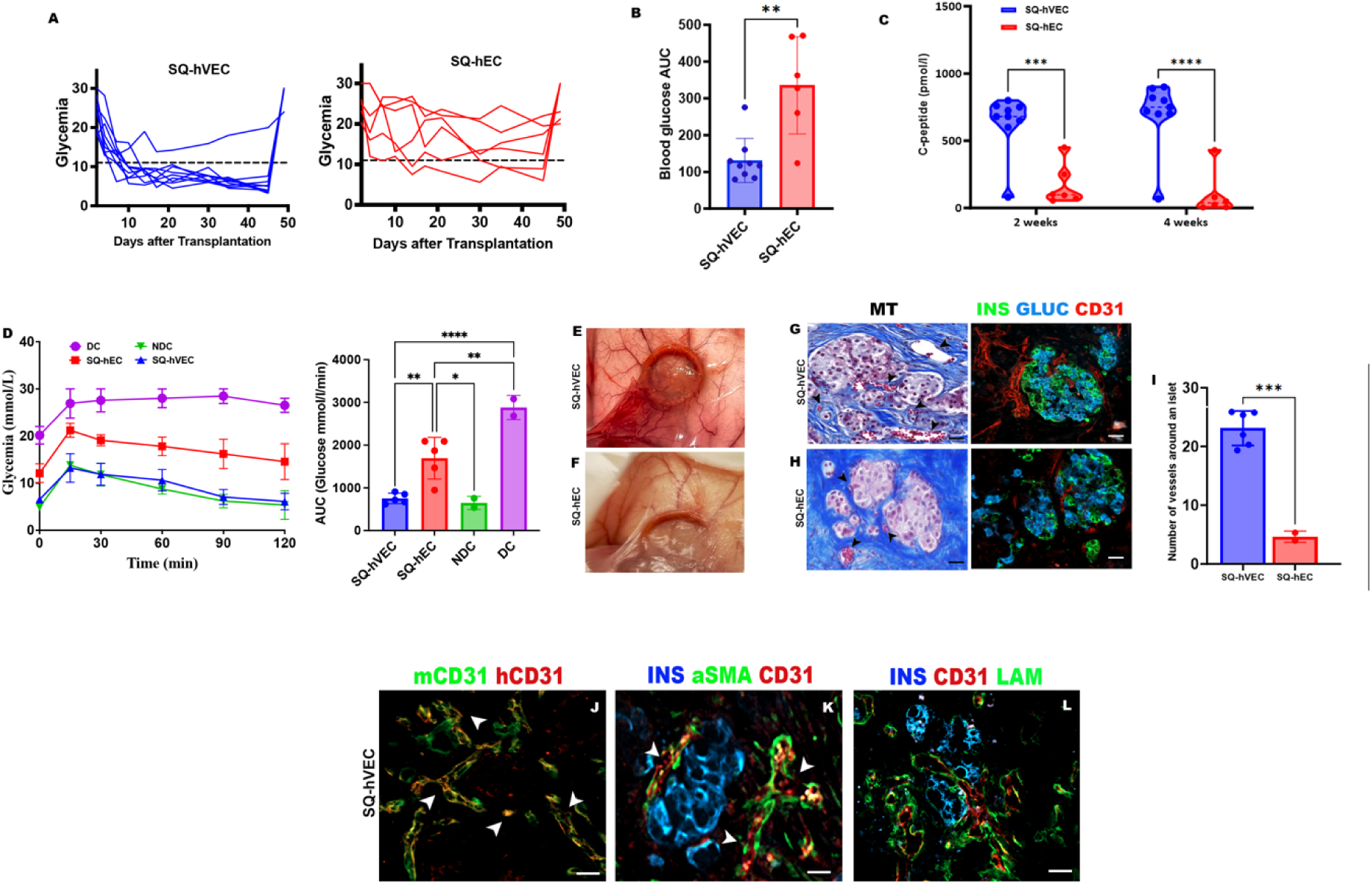
Performance of human VECs after subcutaneous transplantation in diabetic mice. (A) Blood glucose measurements following transplantation. Data are presented as mean ± SEM, one-way ANOVA, SQ-hVEC vs. SQ-hEC, ****p < 0.0001; (B) AUC of blood glucose levels from day −7 to day 30. Data are presented as mean ± SD, Unpaired t-test, p**=0.0012. (C) IPGTT results for SQ-hVEC (n = 5), SQ-hEC (n = 5), diabetic control (DC, n = 2), and non-diabetic control (NDC, n = 2). Data are presented as mean ± SEM. (D) AUC analysis of IPGTT. Data are presented as mean ± SD, one-way ANOVA with Tukey’s correction, SQ-hVEC vs. SQ-hEC., ** p =0.0057, SQ-hVEC vs. NDC, n.s., p=0.9803, SQ-rVEC vs. DC, p****<0.0001, SQ-hEC vs. NDC, p*=0.0168, SQ-hEC vs. DC, p**=0.0077 (E,F) Explanted constructs 45 days after transplant. (G, H) Cross-sectional images of retrieved constructs on day 45 showing insulin/glucagon-positive islets and vascularization of the graft. Masson’s trichrome staining and fluorescence immunohistochemistry with CD31 (red) revealed blood vessels containing RBCs (black arrowheads (I) Blood vessel density in retrieved SQ-hVEC and SQ-hEC. Data are presented as mean ± SD, two-tailed unpaired t-test, ***p=0.0002. (J-L) Immunofluorescent staining of SQ-hVECs for insulin (blue), human CD31 (red), mouse CD31 (green), α-SMA (green), and laminin (green). Chimeric blood vessels and mature vessels containing RBCs are indicated by white arrowheads. Scale bars, 20 μm.

Explanted constructs showed clear differences between the two experimental groups. SQ-hVECs were fully integrated with surrounding tissues and displayed well-defined, blood-perfused vascular structures of various calibers surrounding the graft (Figure 8E). In contrast, SQ-hECs displayed reduced vascularization and less effective integration with host tissues (Figure 8F). Histological analysis showed viable islets with normal morphology in both groups (Figure 8G, H). However, islets in the SQ-hVEC group were surrounded by significantly more blood vessels than those in the SQ-hEC group, consistent with the observed improvements in diabetes correction (Figure 8I). Immunostaining for human and mouse CD31 demonstrated the presence of chimeric blood vessels, indicating anastomoses and vascular remodeling (Figure 8J). Functional maturity of the vasculature was confirmed by vessels with lumens containing erythrocytes, positively stained for human CD31 and α-SMA (Figure 8K). Additionally, islets in the SQ-hVEC group regained laminin expression, an important extracellular matrix protein primarily deposited by endothelial cells (Figure 8L). Laminin was observed both within the islets and in the surrounding tissue.

These findings demonstrate that prevascularizing Amniogel-based endocrine constructs significantly enhances islet engraftment and function in subcutaneous transplantation sites, leading to superior glycemic control.

## DISCUSSION

Recapitulating the key elements of the pancreatic niche, specificaly the ECM microenvironment and vascular network is fundamental requirement for bioengineering a functional endocrine pancreas (*50, 51*). While, collagen-based hydrogels and decellularized ECM matrices support cellular integration and vascularization (*19, 30, 31, 52–58*), the scarcity of human ECM (*59, 60*), its compromised structural integrity and cytotoxic residues associated with conventional detergent-based decellularization techniques (*61, 62*) hinder clinical translation of these strategies.

In this study, we developed, Amniogel, a human amniotic membrane-derived hydrogel, that meets these requirements. By mimicking the ECM and supporting vascularization, Amniogel recreates an islet-like microenvironment. Our bioengineered endocrine constructs incorporate a collagen-rich, biocompatible matrix reminiscent of the native islet niche (*63, 64*), and built-in dense, stable capillary network directly interfaced with the encapsulated islets, enabling efficient glucose sensing and insulin release even in the traditionally poor subcutaneous site.

From a translational perspective, the use of Amniogel platform offers several key advantages. Amniotic membrane is an ethically sourced, established, FDA-approved clinical-grade material widely used in in wound healing and ophthalmology (*65–69*). Our GMP-compliant manufacturing protocol is scalable and reproducible. All reagents used in Amniogel production are available in clinical-grade formulations. Additionally, its cost-effective production provides a viable alternative to expensive purified ECM components. Beyond its biochemical composition, Amniogel demonstrates optimal self-assembly, structural integrity supporting cell adhesion and proliferation, and robust biocompatibility in immunocompetent hosts. Utilization of the host-derived endothelial for vascularization is another important factor to facilitate clinical translation. Taken together, these features give Amniogel a significant regulatory advantage by capitalizing on existing approval frameworks rather than requiring entirely new pathways (*70*).

A central finding of our work is that providing an appropriate human ECM substrate significantly boosts islet graft health by engaging cell–matrix survival pathways. Isolation procedures often disrupt the islet ECM, triggering β-cell apoptosis via β1-integrin signaling (*71*). Previous studies have demonstrated supplementing ECM proteins improves islet survival and function (*72*). In line with these observations, our results showed that islets embedded in Amniogel exhibited improved glucose-responsive insulin secretion and increased expression of β1-integrin, a key mediator of cell-ECM adhesion critical for β-cell survival. Additionally, Amniogel-embedded islets displayed reduced pro-apoptotic CASP3 and elevated anti-apoptotic BCL2 expression. These findings suggest that restoring cell–matrix interactions effectively reduce β-cell apoptosis, consistent with the established role of β1-integrin signaling in promoting β-cell survival (*64, 73, 74*). Furthermore, we observed enhanced intercellular communication in generated VECs, via connexin-mediated gap junctions. Cx-36, predominantly expressed in β-cells and facilitates coordinated insulin release (*37, 75*) while, Cx-43 mediates endothelial–β-cell communication, increasing islet size and insulin content (*75–77*). Our results show significant upregulation *GJD2* (5.27-fold in Amniogel; 8.08-fold in vascularized constructs) and *GJA1* (6.3-fold in Amniogel; 11.4-fold in vascularized constructs) compared to suspension cultures, suggesting that matrix components enhance gap-junctional intercellular communication. However, it remains unclear whether this effect is specifically due to laminin or broader ECM-mediated mechanisms that stabilize islets. ECM– integrin interactions strongly influence connexin expression and intercellular communication in other cell types (*78, 79*), suggesting a similar mechanism in our observations. Given laminin’s known role in integrin activation and connexin regulation (*80–82*), β1-integrin signaling may promote ECM-mediated gap junction formation, thus enhancing synchronized insulin secretion in β-cells. These molecular changes were associated with decreased basal insulin secretion and improved glucose responsiveness in vascularized constructs, further supporting the ECM’s functional relevance in β-cell physiology. Additional studies are required to elucidate the precise contribution of laminin–integrin signaling to gap junction assembly and to clarify whether these effects arise primarily from BOEC interactions, β1-integrin signaling, or both.

The development of functional 3D vascular networks remains a critical hurdle in tissue engineering (*83*). Amniogel promotes endothelial cell self-assembly into interconnected vascular structures, creating a microenvironment conducive to sustained nutrient and oxygen delivery to encapsulated islets. This spatial arrangement ensures insulin secretion occurs near blood vessels, a prerequisite for optimal graft function in poorly vascularized transplant sites. Furthermore, utilization of BOECs, which can be derived from the recipients themselves is a key advantage of this approach. Evidence shows that humoral immune responses significantly contribute to vascularized allograft failure (*84, 85*), as donor-specific antibodies (DSA) primarily target the mismatched HLA molecules, expressed on graft endothelial cells (*86*). Consequently, antibody-mediated rejection (AMR) predominantly affects graft microvasculature (*87*). Experimental evidence further shows that endothelial chimerism—where recipient-derived vessels vascularize the graft—confers resistance to AMR (*86*). Moreover, recent studies indicate allogeneic graft endothelia are also susceptible to innate immune attacks by recipient natural killer (NK) cells, causing antibody-independent graft injury (*88–90*). Thus, replacing donor endothelial cells with autologous BOECs represents a promising strategy to reduce immune activation and prolong graft survival. Importantly, BOECs from both diabetic and non-diabetic patients have been shown to enhance islet engraftment with equal efficacy (*91*), reinforcing their potential for clinical use. Their accessibility, robust proliferative capacity, and intrinsic ability to form functional vascular networks render them highly suitable for improving engineered tissue stability and function (*92, 93*). Our findings further demonstrate that BOEC-derived vasculature surrounding islets produces a basement membrane-like ECM enriched in laminin, mimicking the native islet niche. This observation aligns with previous reports that β-cells rely on intra-islet endothelial cells to synthesize basement membrane constituents (*94*).

In vivo studies confirmed that both ECM and prevascularization are essential for optimal engraftment of the engineered constructs. In a marginal islet transplant model within the highly vascularized epididymal fat pad, Amniogel-embedded islets exhibited significantly improved engraftment compared to islets transplanted alone. However, non-vascularized constructs at the same islet dose failed to restore normoglycemia, likely due to insufficient oxygenation and nutrient exchange in this poorly vascularized site. These findings align with previous reports indicating that although ECM integration enhances cell survival, robust vascularization remains critical for effective glucose sensing and insulin secretion, particularly in the subcutaneous space, which remains challenging despite clinical benefits [33, 93]. This limitation was overcome by preveascularizing subcutaneously implanted endocrine constructs, resulting in significantly improved islet survival and function. Notably, 96% of SQ-rVEC recipients achieved normoglycemia within one week, compared to 20% in the SQ-rEC group and none in the SQ-rI group. Histological analysis confirmed increased vessel density in SQ-rVEC grafts, reinforcing the crucial role of vascularization for successful engraftment. Replication of these experiments with human islets yielded similar results, including rapid elevation of circulating human C-peptide post-transplantation. Explanted human grafts displayed excellent islet morphology, hormone expression and extensive vascularization. The presence of red blood cells and α-SMA-positive vessels indicated functional maturity, while chimeric vessels and restored laminin expression further confirmed successful vascular integration.

Taken together, these results demonstrate that vascularized endocrine constructs effectively address the critical limitations associated with subcutaneous islet transplantation by facilitating rapid vascular integration and sustaining islet function.

Beyond these advantages, the versatility of Amniogel opens opportunities for broader application in regenerative medicine. the versatility of Amniogel broadens its potential in regenerative medicine, making it suitable for stem cell-derived insulin-producing cells, xenogeneic islets, or other therapeutic cell types. Additionally, Amniogel could be modified with immunomodulatory cells or molecules to optimize the local immune microenvironment and enhance engraftment outcomes. Nevertheless, any modifications must be carefully optimized to maintain the established safety profile of amnion-derived products and avoid introducing additional regulatory complexities (*95*).

## MATERIALS AND METHODS

### Study Design

The objective of this study was to develop a clinically scalable, vascularized endocrine construct (VEC) using a human amniotic membrane–derived hydrogel for β-cell replacement therapy. Amniogel was manufactured under GMP-compatible conditions and evaluated for ECM content, gelation properties, and biocompatibility. Pancreatic islets, alone or co-encapsulated with blood outgrowth endothelial cells (BOECs), were embedded in Amniogel and assessed for viability, vascular network formation, and insulin secretion in vitro.

Therapeutic efficacy was tested in streptozotocin-induced diabetic NSG mice following transplantation of constructs into the epididymal fat pad or subcutaneous space. Functional outcomes included glycemic control, glucose tolerance, and histological evidence of engraftment and vascular integration.

Sample size was determined using power analysis (G*Power). Randomization was performed after diabetes induction, and group allocation was concealed during transplantation. Investigators assessing outcomes were blinded to group assignments when feasible, and were independent from those conducting the experiments.

### Amniogel Preparation

To generate Amniogel in line with GMP for future clinical use, we formulated a precise five-step protocol. The process includes the harvesting of tissue, decellularization, sterilization, freeze-drying, mincing, digestion, purification and neutralization (Fig. 1A).

Immediately after delivery, placenta was immersed in sterile PBS supplemented with antibiotics (100 U/ml penicillin, 100 mg/ml streptomycin, and 0.25 mg/ml amphotericin B) and transferred to the laboratory. Under sterile conditions, the human amniotic membrane (hAM) was separated, rinsed to remove blood clots, and decellularized using 10X TrypLE™ Select (ThermoFisher Scientific) for 40 minutes at 37°C with shaking. Decellularized membranes were washed, freeze-dried, minced, and digested with 1 mg/mL porcine pepsin (Sigma Aldrich) in 0.04 N HCl for 48 hours at room temperature. The solubilized hAM, rreferred to as “pre-gel” was centrifuged, adjusted to 10 mg/mL, aliquoted, and stored at −80°C.

Gelation was achieved by neutralizing pre-gels (pH 7.4) with MCDB 131 medium and NaHCO₃, yielding 8 mg/mL Amniogel. Neutralized gel (62%) was mixed with cell suspension (38%), yielding hydrogels with 5 mg/mL matrix proteins, cast into silicon molds or droplets, and polymerized at 37°C for 30 minutes. Amniogel batches were assessed for DNA/protein content, gel formation, and biocompatibility.

### Evaluation of Amniogel’s Impact on Islet Function

Rat or human islets (1000 IEQ) were suspended in 100 μL of neutralized Amniogel, loaded into silicone molds, and incubated at 37 °C to facilitate gelation. The gelled constructs were cultured for 7 days, with the medium replaced every other day. Suspension-cultured islets served as experimental controls.

To evaluate the potential cytoprotective effects of Amniogel on islet cells under ischemic conditions, an experiment was conducted with two groups: islet cells embedded within Amniogel and control islet cells cultured without the gel. Both groups were exposed to a 16-hour hypoxic environment to simulate ischemia.

The experiment was performed in a humidified incubator connected to a nitrogen supply, maintaining a gas mixture of 1% oxygen, 5% CO₂, and 94% nitrogen. This setup accurately replicated the low-oxygen environment characteristic of ischemia. The incubator temperature was consistently maintained at 37 °C throughout the experiment.

Static glucose-stimulated insulin secretion (GSIS) assays were conducted to assess function of islets. To this end, islets embedded in Amniogels or maintained in suspension culture were sequentially incubated in 2.8 mM glucose (basal) and 16.7 mM glucose (stimulated) for 1 h each, and supernatants were collected for insulin quantification. Human islets underwent an additional stimulation step using 5 mM theophylline to enhance insulin secretion. Total insulin content was extracted using acid-ethanol, and the stimulation index (SI) (stimulated/basal insulin secretion) was calculated. All assays were performed in duplicate with 100 IEQ per condition.

### Vascular network formation in Amniogel constructs

GFP-labeled BOECs (2×10⁶/mL) were mixed with Amniogel, dispensed into 10–15 µL droplets, and incubated at 37°C with 5% CO₂ for 30 minutes. Gelled droplets were cultured in vasculogenic media (VM) composed of complete EM supplemented with 40 ng/mL VEGF-A and 20 ng/mL FGF-2 for 7–14 days.

### Assembly of vascularized endocrine constructs

To study vascular network formation and islet vascularization, isolated islets (1000 IEQ) and BOECs (2×10⁶) were encapsulated in 100 μL of Amniogel. The mixture was cast into silicone molds to form 10 μL droplets, polymerized, and cultured at 37 °C for 7 days in optimized vasculogenic medium (OVM), a 1:1 mixture of complete DMEM and VM specifically formulated to maintain islet and endothelial cell viability while promoting vascular network self-assembly. Controls included free islets and islet-laden Amniogels cultured under the same conditions.

### Live Visualization

To study endothelial self-organization and network formation, live microscopy was conducted using an epifluorescent microscope (DMi8 manual microscope; Leica Microsystems, Heerbrugg, Switzerland). This was done without additional staining, as the endothelial cells were labeled with GFP. Observations were initiated 24 hours after casting the construct, allowing sufficient time for polymerization and stabilization to occur within the construct.

### Lumen Formation Analysis and Immunofluorescence

Free-floating constructs were fixed in 4% paraformaldehyde at room temperature for 2 hours and blocked with 3% FBS, 1% BSA, 0.5% Triton X-100, and 0.5% Tween for 2 hours. Primary antibodies against insulin (1:50, DakoCytomation) and CD31 (1:100, DAKO) were applied overnight at 4°C, followed by secondary antibodies (donkey anti-guinea pig Alexa Fluor 488 and donkey anti-mouse Alexa Fluor 594, both 1:500, Jackson ImmunoResearch) for 45 minutes and three PBS washes.

To confirm lumen formation, constructs were incubated with 0.5 mg/mL Texas red-labeled Dextran (10,000 MW, ThermoFisher) in MCDB 131 medium at 37°C and 5% CO₂ overnight, fixed in 4% PFA for 30 minutes, and washed with PBS.

Samples were cleared with RapiClear overnight at room temperature and mounted in iSpacer microchambers. Fluorescent imaging was performed using a Nikon A1R confocal microscope, and Z-stacks were generated with NIS-Elements Imaging Software.

### Implantation of Endocrine Constructs into the Epididymal Fat Pad of Diabetic Mice

Immediately after isolation, 250 rat IEQs were aliquoted into 38 μl of medium, combined with 62 μl of neutralized Amniogel, cast into 10 μL silicone elastomer molds, and polymerized at 37°C. Constructs were cultured for 24 hours. On the day of transplantation, a 0.7 cm incision was made in the peritoneal wall near the genital area to expose the EFP. The islet-laden Amniogels were positioned on the fat pad, folded, and reinserted into the peritoneal cavity. Each mouse received four endocrine constructs containing approximately 250 IEQ (EFP-rEC; n=8). Control mice were transplanted with 250 IEQ alone (EFP-rI; n=8).

### Subcutaneous transplantation of vascularized endocrine constructs

For subcutaneous transplantation, rat or human islets (1000 IEQ) were combined with BOECs (0.2 × 10⁶) and mixed with Amniogel. For human islets, 100 μL of the mixture was cast into 9 mm silicone molds; for rat islets, 10–15 μL droplets were dispensed into silicone elastomer molds. The polymerized constructs were cultured at 37°C in OVM overnight.

Next day, mice were anesthetized with 2.5% isoflurane, and a 0.5 cm incision was made to create subcutaneous pockets. A silicone ring was placed on the muscle and secured with Tisseel fibrin sealant (Baxter AG) to prevent migration. Constructs were inserted into the ring, and the incision was sealed with Tisseel fibrin sealant and sutured using 5–0 silk.

For rat islet constructs (SQ-rVEC), mice (n=18) received 8–10 microgels containing 1000 islets on the left side. Controls included non-vascularized endocrine constructs (SQ-rEC) with identical islet quantities (n=15) and subcutaneously implanted naked islets (n=6).

For human islet constructs (SQ-hVEC; n=9), a single construct. (9mm diameter) containing 1000 IEQ was implanted on each side (2000 IEQ total). Mice transplanted with nonvascularized constructs (SQ-hEC; n=6), containing equal number of islets served as controls.

### Histopathological analysis and immunostaining

Explanted grafts were fixed in 4% paraformaldehyde and either paraffin-embedded or snap-frozen in OCT compound (Tissue-Tek, Sakura Finetek). Serial 5 µm sections (10 per graft) were prepared for hematoxylin-eosin (H&E), Mason Trichrome (MT), or immunofluorescence staining.

For immunostaining, sections were permeabilized, blocked, and incubated overnight with primary antibodies against insulin (1:50, DakoCytomation), glucagon (1:4000, Sigma-Aldrich), human-specific CD31 (1:50, DakoCytomation), CD31 (1:50, Abcam), CD34 (1:2000, Abcam), laminin (1:30, Sigma-Aldrich), collagen IV (1:30, Bio-Rad), CD45 (1:100, Abcam), and CD11b (1:200, Abcam). Secondary antibodies included goat anti-mouse, anti-rabbit, anti-guinea pig (1:300, ThermoFisher Scientific), goat anti-guinea pig (1:500, Jackson ImmunoResearch), donkey anti-rabbit Alexa Fluor 594 (1:500, Jackson ImmunoResearch), and rat anti-mouse Alexa Fluor 594 (1:200, Abcam). Sections were mounted with DAPI-containing mounting medium (ProTaqs MountFluor Anti-Fading).

Images were captured using a Zeiss Axioscan.Z1 slide scanner, and morphometric and fluorescence analyses were performed with ImageJ software.

In H&E and MT-stained images, blood vessels were identified by luminal structures containing erythrocytes. Vessel numbers within grafts were quantified, and vessel/graft and vessel/β-cell ratios were calculated as percentages of graft and insulin-positive areas. Human vasculature was confirmed by erythrocyte-containing vessels positive for human CD31 and α-SMA staining.

## Statistical analysis

Statistical analyses were performed using GraphPad Prism 10.0. All experiments were repeated at least three times. Specific statistical tests are detailed in the figure legends, with significance defined as P < 0.05.

## List of Supplementary Materials

Materials and Methods

Fig. S1 or Fig S9

Table S1

References (*1,2*)

## Author contributions

E.B. conceptualized and supervised the project.; E.B. and K.B. developed the methodology; K.B., F.L. and M.H. performed animal breeding and conducted experiments; K.B., F.L., M.H., R.H., J.B., A.M., L.M.F., and F.C. performed analysis of results; K.B., and M.H. performed visualization.; A.F. and A.C. produced BOECs; E.B. K.B. and F.L. wrote the original draft.; A.F., L.P., O.T., J.S, M.C. and E.B. contributed to the design of the experiments.; A.F., C.O., J.B., A.M., L.M.F., M.C., B.MT., L.P., A.C., O.T., J.S., M.H., L.WB., P.C. and VANGUARD consortium critically reviewed and edited the paper.

## Data and materials availability

All data are available in the main text or the supplementary materials.

## § Members of the VANGUARD consortium are as follows

Thierry Berney, Nicerine Krause, Shota Tsikhiseli, University of Geneva, Department of Surgery, Geneva, Switzerland

Alessia Cucci, Chiara Borsotti, Simone Assanelli, Department of Health Sciences, University of Piemonte Orientale, Novara, Italy

Cataldo Pignatelli, IRCCS Ospedale San Raffaele, Diabetes Research Institute, Milano, Italy

Emma Massey, Dide de Jongh, Erasmus MC, University Medical Centre Rotterdam, The Netherlands

Eline Bunnik, Dept. of Medical Ethics, Philosophy and History of Medicine, University Medical Centre Rotterdam, The Netherlands

Antonia J. Cronin, King’s College, London, UK

Devi Mey, Chiara Parisotto European Society for Organ Transplantation, Padova, Italy

Patrick Kugelmeier, Petra Wolint, Kugelmeiers AG, Erlenbach, Switzerland

Margaryta Schaltegger, Julia Götz, Jeanette Müller, Accelopment Switzerland Ltd.

