## Supplementary Material and Figures for "Bioengineering of the implantable vascularized endocrine constructs for insulin delivery suitable for clinical upscaling"

**Supplementary Information**

### MATERIALS AND METHODS

#### Animals

Male, 6–8-week-old NOD.Cg-Prkdc$^{scid}$ Il2rg$^{tm1Wjl}$/SzJ (NSG), C57BL/6 mice and male 8-week-old Sprague-Dawley (SD) rats (250–300 g) were bred in house at the University of Geneva with free access to food and water. All animal procedures were approved by the University of Geneva Institutional Animal Care and Use Committee.

SD rats were used as islet donors. Pancreatic islets were isolated by injecting ice-cold collagenase type V into the pancreatic duct. The pancreas was excised, digested at 37°C, and the islets were purified using a discontinuous Ficoll gradient. Isolated islets were cultured in DMEM supplemented with 10% Fetal Bovine serum (FBS), 1 mmol/L sodium pyruvate, 11 mmol/L glucose, 0.05 mmol/L 2-mercaptoethanol, 2 mmol/L L-glutamine, 100 U/mL penicillin, and 0.1 mg/mL streptomycin.

#### Human samples and cells

Studies involving human tissues were approved by the Commission Cantonale d'Ethique de la Recherche (CCER), in compliance with the Swiss Human Research Act (810.30). Placental tissues, de-identified and discarded, were obtained from consenting women undergoing elective cesarean section, under CCER protocol PB_2017-00101 (14-273).

Human islets isolated from deceased multiorgan donors. The use of human islets for research was approved by CCER protocol 2016-01979. Donor information is listed in Supplementary table 1. In this study, islets from 8 donors were used.

Purified islets were cultured in CMRL medium containing containing 5.6 mmol/L glucose and supplemented with 2 mmol/L L-glutamine, 100 U/ml Penicillin and 0.1 mg/ml Streptomycin, 25 mmol/l HEPES and 10% FBS in a 5% $CO_2$ incubator.

Blood outgrowth endothelial cells (BOECS) have been isolated from human peripheral blood as previously described (*1*) and expanded up to passage 12. Primary BOECs were cultured in Collagen Type 1-coated tissue culture flasks in MCDB 131 medium containing proprietary supplements. For GFP expression BOECs were plated at a $10^4$ cells/cm$^2$ density and after 6 h transduced with a lentiviral vector carrying the GFP under the control of the ubiquitous promoter of the Phosphoglycerate kinase (LV-

PGK.GFP) using a multiplicity of infection (MOI) of 10. After 16 h incubation, fresh medium was added and, 72 h later, half of the cells were harvested for subsequent analysis, while the other half was further cultured for expansion.

**Lentiviral vector production**

Third generation self-inactivating LVs were produced as published previously (*2*). Briefly, 293T cells were expanded in IMDM, 10/ FBS and transiently co-transfected by the calcium phosphate precipitation method with third-generation packaging plasmids : pMDLg/pol and pRSV-Rev, with the envelope construct (pMD.VSV.G), and the transfer vector construct (pPGK.GFP.LV). The cell supernatant was harvested after 36 hours, and LV particles were concentrated by ultracentrifugation. LV titers were determined on 293T by qPCR .

**DNA quantification and biochemical characterization**

To assess DNA content in native and decellularized hAM, tissue samples were digested with Proteinase K at 55°C overnight. DNA was isolated using saturated 6 M NaCl, precipitated with isopropanol, washed with 70% ethanol, and resuspended in RNase/DNase-free water. DNA concentration and purity were measured spectrophotometrically (NanoDrop 2000c, Thermo Scientific) using absorbance at 260 nm and 280 nm.

Biochemical composition of Amniogel batches was evaluated by quantifying collagen and glycosaminoglycans using the QuickZyme Hydroxyproline Kit and Blyscan™ assay (Biocolor), respectively, following the manufacturers' protocols.

**Protein identification by ESI-LC-MSMS**

Protein composition of Amniogel (n=3) was analyzed by mass spectrometry. Samples were lysed in RIPA buffer, centrifuged, and protein pellets resuspended in thiourea/urea lysis buffer. Following overnight acetone precipitation, pellets were treated with urea, ammonium bicarbonate, dithioerythritol, and iodoacetamide, then digested with trypsin. Supernatants were desalted, dried, and stored at -20°C.

Mass spectrometry analysis was performed at the Proteomics Core Facility, University of Geneva, using a Q-Exactive Plus Hybrid Quadrupole-Orbitrap Mass Spectrometer coupled with an Easy nLC 1000 system. Peptides were separated using specialized columns over a 90-minute gradient. MS survey scans and data-dependent analysis were optimized with specific parameters.

Data was processed using MS Convert from ProteoWizard and searched against the Human Reference Proteome and an in-house contaminant database with Mascot. Identifications were validated in Scaffold 4.8.4 based on probability and False Discovery Rate (FDR), with protein annotations obtained from NCBI GO terms.

**Scanning Electron Microscopy and Transmission Electron Microscopy**

The morphology of decellularized hAM and Amniogel was examined using scanning electron microscopy (SEM) and transmission electron microscopy (TEM). For SEM, samples were fixed in 2.5% glutaraldehyde, washed in 0.1 M phosphate buffer (pH 7.4), post-fixed with 1% $OsO_4$ and 1.5% potassium ferrocyanide, dehydrated through graded ethanol, and dried using a critical point dryer (Leica EM CPD030). Samples were sputter-coated and imaged with a JOEL JSM-6510LV SEM at 10,000× magnification. Mean pore sizes were calculated from 138 measurements across six random areas per sample (n=3).

TEM analysis of collagen fiber ultrastructure utilized glow-discharged, carbon-coated Quantifoil grids incubated with 3 μL samples. Grids were washed, stained with 2% uranyl acetate, and imaged using a ThermoFisher G20 Sphera microscope at 120 kV with an FEI Eagle detector (4k × 4k resolution). Imaging parameters included a defocus range of -1 to -2 μm, controlled with SerialEM software. Micrographs were aligned using IMOD (Etomo module), and collagen fiber measurements were quantified with Fiji software.

**Turbidimetric Gelation Kinetics**

Gelation kinetics were analyzed using a SpectraMax spectrophotometer (405 nm). Solutions (5 mg/mL) were stored at 4°C, transferred to a chilled 96-well plate (100 μL/well, in triplicate), and maintained on ice. The plate was then placed in a pre-warmed spectrophotometer at 37°C, and turbidity measurements were recorded every 2 minutes for 1.5 hours. Three independent tests (n = 3) were conducted on a single batch of rat tail collagen type I (Sigma Aldrich), and three additional tests (n = 3) were performed on separate Amniogel batches, with each test conducted in triplicate. Parameters measured included t1/2 (time to half-maximal turbidity), tlag (lag phase determined from the linear portion of the curve), and gelation speed (maximum slope of the growth phase).

**Oscillatory Rheology**

The rheological properties of Amniogel were assessed using a modular dynamic rheometer (Haake Mars 40, ThermoFisher Scientific). Both uncrosslinked (as-prepared) and crosslinked (gelled) samples were analyzed. Measurements for uncrosslinked samples were conducted at 4°C, while crosslinked samples were tested at 37°C to simulate in vitro and in vivo conditions. All experiments were performed in triplicate.

For rotational tests, 400 µL of sample was loaded onto a parallel plate geometry (0.096 mm gap) pre-set to the desired temperature, and shear rates from 0 to 200 $s^{-1}$ were applied.

Oscillatory tests were performed on crosslinked samples to evaluate viscoelastic properties, as uncrosslinked samples exhibited liquid-like behavior. Samples were subjected to oscillatory frequencies from 1 to 10 Hz, with five measurements per step.

**Contraction of Amniogels**

The contraction of cell-laden Amniogels was evaluated using microscopic imaging. Amniogels were prepared by incorporating GFP-labeled BOECs at cell densities of $0.5 \times 10^6$ and $2 \times 10^6$ cells/mL into gels with concentrations of 8 mg/mL and 5 mg/mL. A 100 µL volume of each mixture was cast into 9 mm silicone molds and polymerized at 37 °C. The polymerized samples were cultured as free-floating constructs in complete media for seven days in 6-well plates, with media changes performed every other day. Unseeded Amniogels at equivalent concentrations were used as controls.

The top surface areas of the hydrogels were measured at multiple time points using ImageJ software (NIH, Bethesda, MD). The borders of the hydrogels were manually traced, and the surface areas were calculated as a percentage of the initial unseeded hydrogel area to quantify contraction.

**Cytocompatibility Testing**

Primary BOECs ($1\times10^3$ cells/$cm^2$) were seeded on plates coated with Amniogel (0.2 mg/mL) and cultured for 7 days in their respective media. Controls included BOECs seeded on collagen I-treated and tissue culture-treated plates.

Cell viability was evaluated using fluorescein diacetate (FDA) and propidium iodide (PI) staining (Sigma-Aldrich). Fluorescent signals were observed under epifluorescent microscope (DMi8, Leica Microsystems). Viability was quantified as the ratio of live cells to total cells.

**In Vivo Biocompatibility of Amniogel**

To assess in vivo biocompatibility, eight-week-old C57BL/6 mice (n=3; 25 g) were used. Empty Amniogels were cast into 9 mm round silicone molds one day prior to surgery. Mice were anesthetized with 2.5% isoflurane in oxygen, and anesthesia was maintained throughout the procedure. A 0.5 cm incision was made to create a subcutaneous pocket overlying the muscle. A silicone ring (9 mm inner diameter, 1 mm width, 1 mm thickness) was placed on the muscle, and the Amniogel was positioned inside. The incision was sutured with 4–0 silk. Each mouse received one hydrogel in the ventral subcutaneous space (SQ). Grafts were explanted after four weeks, fixed in 4% paraformaldehyde overnight, and processed for histological analysis.

**Establishment of the Diabetes Models**

Male NSG mice were rendered diabetic via a single intraperitoneal injection of streptozotocin (160 mg/kg, Sigma-Aldrich) one week prior to transplantation. Mice with blood glucose levels >28 mmol/L for three consecutive days were selected and randomly assigned to control or treatment groups prior to transplantation.

**Blood Glucose Monitoring and Assessment of Metabolic Function of the Grafts**

Islet graft function was monitored by measuring non-fasting blood glucose (BG) daily during the first week post-transplantation and bi-weekly thereafter using a portable glucometer (Freestyle Precision, Abbott Diabetes Care, Switzerland). Diabetes reversal was defined as two consecutive BG readings below 11.1 mmol/L.

Four weeks post-transplantation, an intraperitoneal glucose tolerance test (IPGTT) was performed. Mice were fasted overnight, injected intraperitoneally with glucose (2.0 g/kg), and BG levels were measured at baseline and 15, 30-, 60-, 90-, and 120-minutes post-injection. Serum C-peptide levels were quantified using an ELISA (Mercodia).

At the end of the experiment, constructs were explanted from normoglycemic mice, and BG levels were monitored for 3 days to confirm the return to hyperglycemia.

**Quantitative Real-Time Polymerase Chain Reaction (qRT-PCR)**

The ReliaPrepTM RNA Tissue Miniprep System (Promega AG) was used to extract total cellular RNA. cDNA was synthesized using the Quanti Fluor RNA System kit (Promega AG). Gene amplification was achieved by RT-PCR using the Takyon No-Rox SYBR Core Kit blue dTTP (Eurogentec). Primers used for amplification were obtained from Microsynth (Balgach) and are listed in table 1. Gene expressions values were normalized to the housekeeping genes (RPLP0) and calculated based on the comparative cycle threshold Ct method (2−ΔCt method).

**Table S1: Pancreas donor information**

| Donor | Sex | Age | BMI (kg/m$^2$) | DCD/DBD | CIT (hrs) | Purity (%) |
|---|---|---|---|---|---|---|
| **Donor 1** | M | 46 | 24.9 | DBD | 7.5 | 88 |
| **Donor 2** | F | 54 | 32.9 | DBD | 7.8 | 90 |
| **Donor 3** | F | 59 | 23.4 | DBD | 8.25 | 60 |
| **Donor 4** | M | 19 | 24.1 | DBD | 5.6 | 54 |
| **Donor 5** | F | 55 | 31.2 | DBD | 9.5 | 78 |
| **Donor 6** | M | 55 | 28.7 | DBD | 10.55 | 54 |
| **Donor 7** | F | 58 | 28.3 | DBD | 9.5 | 80 |
| **Donor 8** | F | 58 | 21.4 | DBD | 5.05 | 56 |

**Pancreas donor information. (M = male, F = female, BMI = body mass index, DCD = donation by cardiac death, DBD = donation by brain death, CIT = cold ischemia time)**

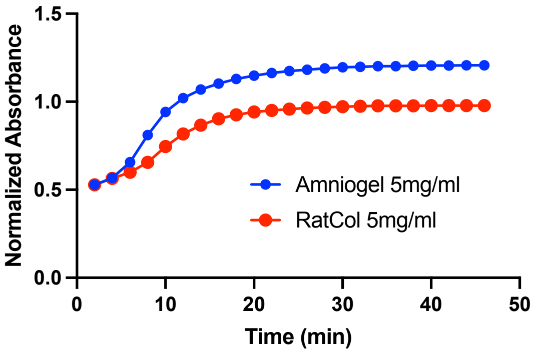


**Fig. S1** Turbidimetric gelation kinetics of collagen I and Amniogel by temperature and time sweep. Average normalized absorbance of gels (n=3 biological replicates) measured at 405 nm shows gelation kinetics over 60 min.

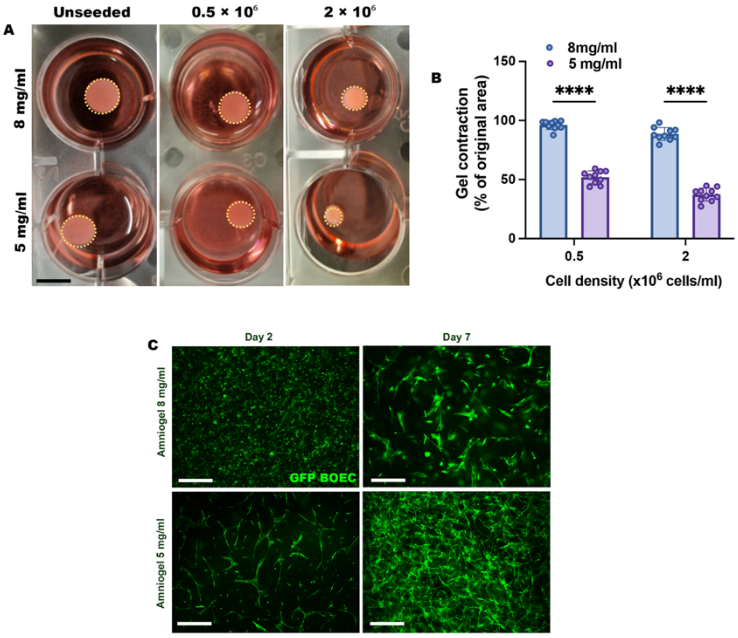


**Fig. S2 Cell mediated contraction of Amniogel.** (A) Representative inverted bright-field images of 8 mg/mL and 5 mg/mL Amniogels with ECM concentrations of 8 mg/mL and 5 mg/mL, seeded with varying LV-GFP transduced BOEC densities, after 7 days of culture. Hydrogel borders are delineated with dotted yellow lines. Scale bars, 2 mm. (B) Quantification of contraction for 8 mg/mL and 5 mg/mL Amniogels at different seeding densities. Contraction is expressed as a percentage of the area of the unseeded control for each ECM concentration. Data are presented as mean ± SD, two-way ANOVA with Sidak correction, ****p<0.0001 ***p=0.0001 (n =10 for three biological replicates per condition at each time point) (C) Representative fluorescent images of the top view of 3D construct showing GFP-labeled BOECs in 8 mg/mL and 5 mg/mL Amniogel after 2 and 7 days in culture, respectively. Scale bars, 250 μm.

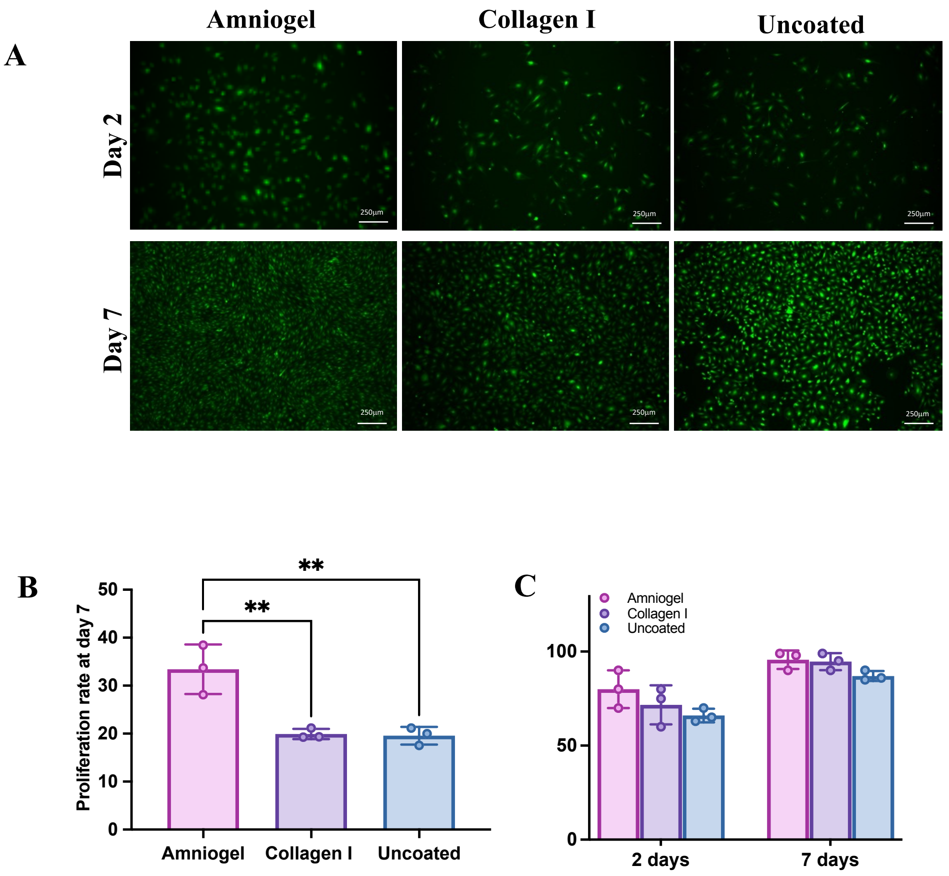


**Fig. S3 Proliferation and viability of BOECs seeded on Amniogel coated plates.** A. GFP labelled BOECs were seeded at 1000 cells/cm$^2$ on either Amniogel, Collagen-1 or tissue culture treated plates. B. Proliferation was calculated by dividing the number of proliferated cells by the number of seeded cells, (n=3), values presented as mean ± SD, one way ANOVA with Tukey's correction, Amniogel vs. Uncoated **p=0.0045;

Amniogel vs Collagen I **p=0.0051 C. Cell viability assay at days 2 and 7. Cells were seeded at 1000 cells/cm$^2$ (n=3), values presented as mean ± SD, no significant difference.

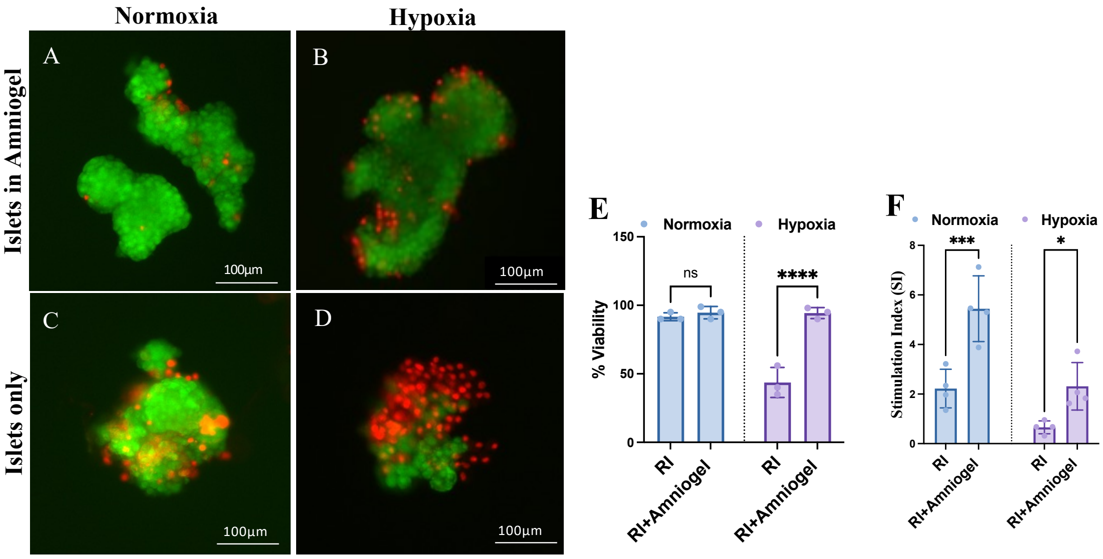


**Figure S4 Amniogel impact on rat islet viability and function under normoxic and hypoxic culture conditions.** A-D. The live (green)/dead (red) staining of rat islets maintained in suspension or in Amniogel under normoxic or hypoxic culture conditions. E. Quantification expressed as percent of live islet area (n = 3 biological replicates), mean ± SD, two-way ANOVA with Šidák correction, ****p <0.0001. F. The GSIS tests of the islets encapsulated in Amniogel and islets in suspension after 24 h culture (n = 4 biological replicates), mean ± SD, two-way ANOVA with Šidák correction, ***p <0.0006, **p <0.0499.

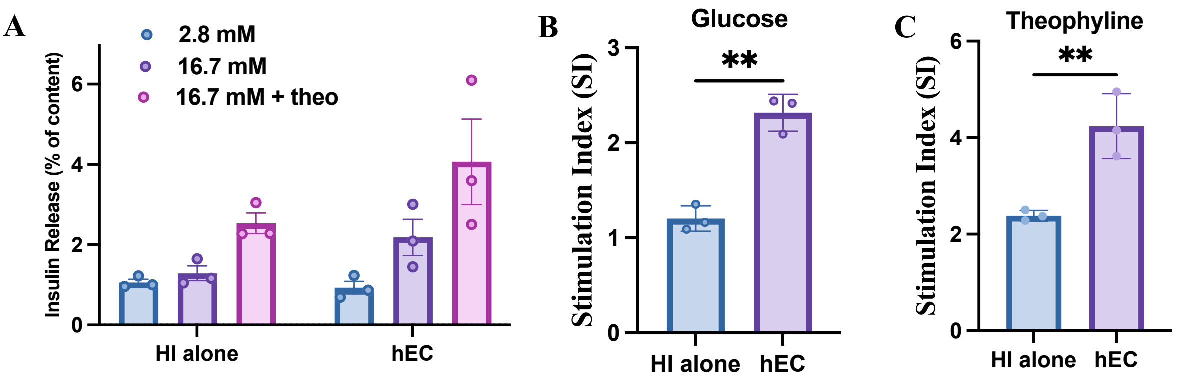


**Fig. S5 Endocrine function of hECs.** (A) Islet function after 7 days of culture was evaluated using GSIS assay with low glucose (2.8 mM), high glucose (16.7 mM), and high glucose plus theophylline (16.7 mM + 5 mM). GSIS data are normalized to the total insulin content. (B, C) Stimulation index (B, high/low glucose) (B, theophylline/low glucose) of hECs or islets in suspension, after 7 days of culture. hECs demonstrate significantly higher stimulation index compared to suspension-cultured islets under both glucose and theophylline stimulation. Data are presented as mean ± SD, two-tailed unpaired t-test, Glucose **p = 0.0012, Theophylline **p = 0.0092 (n = 3 biological replicates).

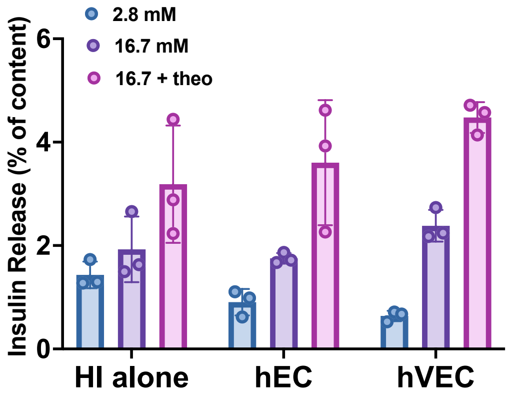


**Fig. S6 Endocrine function of hVECs after 7 days of culture.** GSIS data are normalized to the total insulin content. Vascularized constructs exhibited a superior response to glucose stimulation, characterized by reduced basal insulin secretion, compared to controls (n = 3 biological replicates).

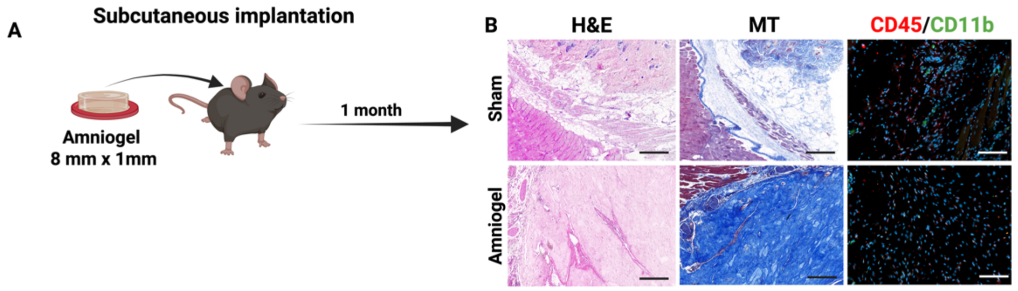


**Fig. S7 Evaluation of in vivo biocompatibility of Amniogel.** (A) Schematic representation of the subcutaneous implantation of Amniogel in C57BL6 mice. (B) Histological analyses 30 days post-implantation: Hematoxylin and Eosin, Masson's Trichrome, CD45 (red), and CD11b (green) staining of cross-sections reveal no significant foreign-body reaction or immune cell infiltration. Scale bars, 200 µm. Created in [https://BioRender.com](https://BioRender.com)

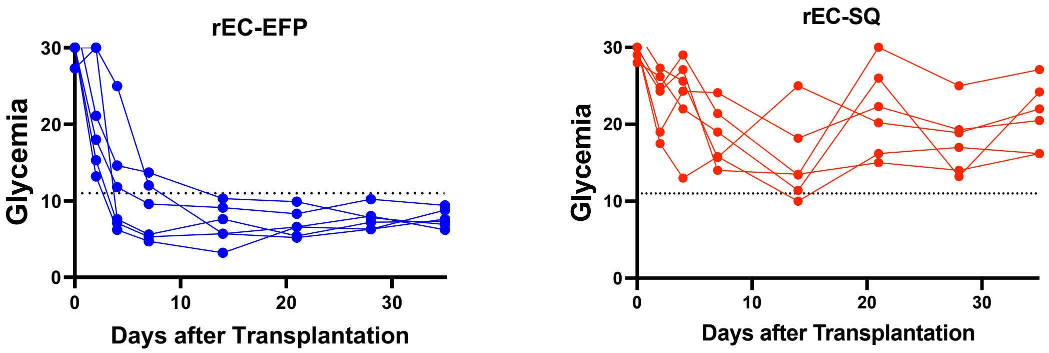


**Fig. S8** Blood Glucose measurements of diabetic NSG mice after transplantation of rECs in EFP and SQ site.

**Refereces:**


1. C. Olgasi, C. Borsotti, S. Merlin, T. Bergmann, P. Bittorf, A.B. Adewoye, N. Wragg, K. Patterson, A. Calabria, F. Benedicenti, A. Cucci, A. Borchiellini, B. Pollio, E. Montini, D.M. Mazzuca, M. Zierau, A. Stolzing, P.M. Toleikis, J. Braspenning, A. Follenzi, Efficient and safe correction of hemophilia A by lentiviral vector-transduced BOECs in an implantable device, Mol Ther Methods Clin Dev 23 (2021) 551-566.
2. A. Follenzi, L. Naldini, Generation of HIV-1 derived lentiviral vectors, Methods Enzymol 346 (2002) 454-65.
